# Variable prediction accuracy of polygenic scores within an ancestry group

**DOI:** 10.1101/629949

**Authors:** Hakhamanesh Mostafavi, Arbel Harpak, Dalton Conley, Jonathan K Pritchard, Molly Przeworski

## Abstract

Fields as diverse as human genetics and sociology are increasingly using polygenic scores based on genome-wide association studies (GWAS) for phenotypic prediction. However, recent work has shown that polygenic scores have limited portability across groups of different genetic ancestries, restricting the contexts in which they can be used reliably and potentially creating serious inequities in future clinical applications. Using the UK Biobank data, we demonstrate that even within a single ancestry group, the prediction accuracy of polygenic scores depends on characteristics such as the age or sex composition of the individuals in which the GWAS and the prediction were conducted, and on the GWAS study design. Our findings highlight both the complexities of interpreting polygenic scores and underappreciated obstacles to their broad use.

## Introduction

Genome-wide association studies (GWAS) have now been conducted for thousands of human complex traits, revealing that the genetic architecture is almost always highly polygenic, i.e., that the bulk of the heritable variation is due to thousands of genetic variants, each with tiny marginal effects (Boyle, Li, and Pritchard 2017; Bulik-Sullivan et al. 2015). These findings often make it difficult to interpret the molecular basis for variation in a trait, but they lend themselves more immediately to another use: phenotypic prediction. Under the assumption that alleles act additively, a “polygenic score” (PGS) can be created by summing the effects of the alleles carried by an individual; this score can then be used to predict that individual’s phenotype (Lynch and Walsh 1998; Gibson 2008; Kathiresan et al. 2008). For highly heritable traits, such scores already provide informative predictions in some contexts: for example, prediction accuracies are 24.4% for height (Yengo et al. 2018) and up to 13% for educational attainment (Lee et al. 2018).

This genomic approach to phenotypic prediction has been rapidly adopted in three distinct fields. In human genetics, PGS have been shown to help identify individuals that are more likely to be at risk of diseases such as breast cancer (e.g., Khera et al. 2018; Inouye et al. 2018; Mavaddat et al. 2019; Khera et al. 2019). Based on these findings, a number of papers have advocated that PGS be adopted in designing clinical studies, and by clinicians as additional risk factors to consider in treating patients (Khera et al. 2018; Torkamani, Wineinger, and Topol 2018). In human evolutionary genetics, several lines of evidence suggest that adaptation may often take the form of shifts in the optimum of a polygenic phenotype and hence act jointly on the many variants that influence the phenotype (Pritchard and Di Rienzo 2010; Berg and Coop 2014; Hoellinger, Pennings, and Hermisson 2019). In this context, PGS are used to examine the evolutionary history of the set of alleles known to impact a complex trait of interest, e.g., height (Berg and Coop 2014; Field et al. 2016; Berg et al. 2019; Uricchio et al. 2019; Edge and Coop 2019; Speidel et al. 2019). Finally, in various disciplines of the social sciences, PGS are increasingly used to distinguish environmental from genetic sources of variability (Conley 2016), as well as to understand how genetic variation among individuals may cause heterogeneous treatment effects when studying how an environmental influence (e.g., a schooling reform) affects an outcome (such as BMI) (Barcellos, Carvalho, and Turley 2018; Davies et al. 2018). In these applications, the premise is that PGS will “port” well across groups—that is that they remain predictive not only in samples very similar to the ones in which the GWAS was conducted, but also in other sets of individuals (henceforth “prediction sets”).

As recent papers have highlighted, however, PGS are not as predictive in individuals whose genetic ancestry differs substantially from the ancestry of individuals in the original GWAS (reviewed in Martin et al. 2019). As one illustration, PGS calculated in the UK Biobank predict phenotypes of individuals sampled in the UK Biobank better than those of individuals sampled in the BioBank Japan Project: for instance, the incremental *R*^2^ for height is approximately 11% in the UK versus 3% in Japan (Martin et al. 2019). Similarly, using PGS based on Europeans and European-Americans, the largest educational attainment GWAS to date (“EA3”) reported an incremental *R*^2^ of 10.6% for European-Americans but only 1.6% for African-Americans (Lee et al. 2018).

To date, such observations have been discussed mainly in terms of population genetic factors that reduce portability (Martin et al. 2017, 2019; Kim et al. 2018; Duncan et al. 2018; Francisco and Bustamante 2018; Sirugo, Williams, and Tishkoff 2019). Notably, GWAS does not pinpoint causal variants, but instead implicates a set of possible causal variants that lie in close physical proximity in the genome. The estimated effect of a given SNP depends on the extent of linkage disequilibrium (LD) with the causal sites (Pritchard and Przeworski 2001; Bulik-Sullivan et al. 2015). Thus, LD differences between populations that arose from their distinct demographic and recombination histories will lead to variation in the prediction accuracy of phenotypes across populations (Rosenberg et al. 2018). Because of their distinct demographic histories, populations also differ in the allele frequencies of causal variants. This problem is particularly acute for alleles that are rare in the population in which the GWAS was conducted but common in the population in which the trait is being predicted. Such variants are likely to have noisy effect size estimates in the estimation sample or may not be included in the PGS at all, and yet they contribute substantially to heritability in the target population. Furthermore, causal loci or effect sizes may differ among populations, for instance if the effect of an allele depends on the genetic background on which it arises (e.g., Adhikari et al. 2019). For all these reasons, we should expect PGS to be less predictive across ancestries.

In practice, given that most individuals (79%) included in current GWAS are of European ancestry (Popejoy and Fullerton 2016; Martin et al. 2019), PGS are systematically more predictive in European-ancestry individuals than among other people. As a consequence, the clinical applications and scientific understanding to be gained from PGS will predominantly and unfairly benefit a small subset of humanity. A number of papers have therefore highlighted the importance of expanding GWAS efforts to include more diverse ancestries (Martin et al. 2018, 2019; Wojcik et al. 2018; Sirugo, Williams, and Tishkoff 2019).

Importantly, factors other than ancestry could also impact the accuracy and portability of PGS. For example, the educational attainment of an individual depends not only on their own genotype, but on the genotypes of their parents, due to nurturing effects (Kong et al. 2018), and of their peers, due to social genetic effects (Domingue et al. 2018), as well as of course on non-genetic factors. Also, traits such as height and educational attainment show strong patterns of assortative mating, which can distort estimated effect sizes in GWAS (Domingue et al. 2014; Robinson et al. 2017; Ruby et al. 2018). To what extent these effects remain the same across cultures and environments is unknown, but if they differ, so will the prediction accuracy. More generally, while we still know little about GxE (genotype-environment interactions) in humans, GxE effects are well-documented in other species—notably in experimental settings—and would further reduce the portability of PGS across environments (Lynch and Walsh 1998; Gibson 2008). In addition, environmental variance could differ between groups, which would change the proportion of the variance in the trait explained by a PGS (i.e., the prediction accuracy) even in the absence of genetic differences or GxE effects. Finally, PGS for some traits may include a component of environmental or cultural confounding associated with population structure (Berg et al. 2019; Sohail et al. 2019; Haworth et al. 2019; Lawson et al. 2019). This source of confounding can increase or decrease prediction accuracy, depending on the structure in the prediction samples.

Given these considerations, it is important to ask to what extent PGS are portable among groups within the same ancestry. To explore this question, we stratified the subset of UK Biobank samples designated as “White British” (WB) according to some of the standard sample characteristics of GWAS studies: the ages of the individuals, their sex, and socio-economic status. We chose to focus on these particular characteristics because they vary widely among GWAS samples depending on sample ascertainment procedures. Furthermore, these characteristics have been shown to influence heritability for some traits in a study of a subset of the UK Biobank (Ge et al. 2017), raising the possibility that these choices also influence prediction accuracy. Indeed, for three example traits, we show that there exist major differences in the prediction accuracy of the PGS among these groups, even though they share highly similar genetic ancestries. For a variety of traits, we further demonstrate that prediction accuracy differs markedly depending on whether the GWAS is conducted in unrelated individuals or in pairs of siblings, even when controlling for the precision of the estimates. This finding is again unexpected under standard GWAS assumptions; it underscores the importance of genetic effects that are included in estimates from some study designs and not others and highlights underappreciated challenges with GWAS-based phenotypic prediction.

At present, it is difficult to fully determine the reasons why we see such variable prediction accuracy across these strata and study designs. Contributing factors probably include indirect genetic effects from relatives, assortative mating, varying levels of environmental variance, GxE interaction effects and perhaps undetected environmental confounding. Nonetheless, our results make clear that the prediction accuracy of PGS can be affected in unpredictable ways by known— and presumably unknown—factors in addition to genetic ancestry.

## Results

### Sample characteristics of the GWAS and prediction set can influence prediction accuracy even within a single ancestry

We examined how PGS for a few example traits port across samples that are of similar genetic ancestry but differ in terms of some common study characteristics, e.g., the male:female ratio (henceforth “sex ratio”), age distribution, or socio-economic status (SES). To this end, we limited our analysis to the largest subset of individuals in the UKB with a relatively homogeneous ancestry: 337,536 unrelated individuals that were characterized by the UKB as “White British” (WB) (Bycroft et al. 2017). In all analyses, we further adjusted for the first 20 principal components of the genotype data, to account for any population structure within this set of individuals (**Materials and Methods**).

In all analyses, we randomly selected a subset of individuals to be the prediction set; we then conducted GWAS using the remaining individuals and built a PGS model by LD-based clumping of the associations (**Materials and Methods**). To examine the reliability of the prediction, we considered the incremental *R*^2^, i.e., the *R*^2^ increment obtained when adding the PGS to a model with only covariates (referred to as “prediction accuracy” henceforth). Whether this measure is appropriate depends on how PGS are to be used; it is not an obvious choice in human genetics, where the goal is often to identify individuals at high risk of developing a particular disease (i.e., in the tail of the polygenic score distribution). Nonetheless, because it has been widely reported in discussions of portability across genetic ancestries (e.g., Lee et al. 2018; Martin et al. 2019), we also used it here.

As a first case, we considered the prediction accuracy of a PGS for diastolic blood pressure in prediction sets stratified by sex, motivated by reports that variation in this trait may arise for somewhat distinct reasons in the two sexes (Reckelhoff 2001; Zhou et al. 2017). We randomly selected males and females as prediction sets (20K individuals each), and used the rest of the individuals for GWAS, matching the numbers of females and males in the GWAS set. Adjusting for mean sex effects and medication use (see Materials and Methods), the prediction accuracy is about 1.31-fold higher for females than for males (Mann-Whitney *p* = 1.4 · 10^-11^; **Fig. 1A**). Thus, despite equal representation of males and females in the GWAS set, the prediction accuracy varies depending on the sex ratio of prediction samples. To examine this further, we repeated the same analysis but performed the GWAS in only one sex. When the GWAS is conducted only in females, the prediction accuracy is about 1.43-fold higher for females than for males; in turn, when GWAS was done in only males, the prediction accuracy in both sexes is similar, as well as somewhat decreased (**Fig. 1B**).

**Figure 1:**
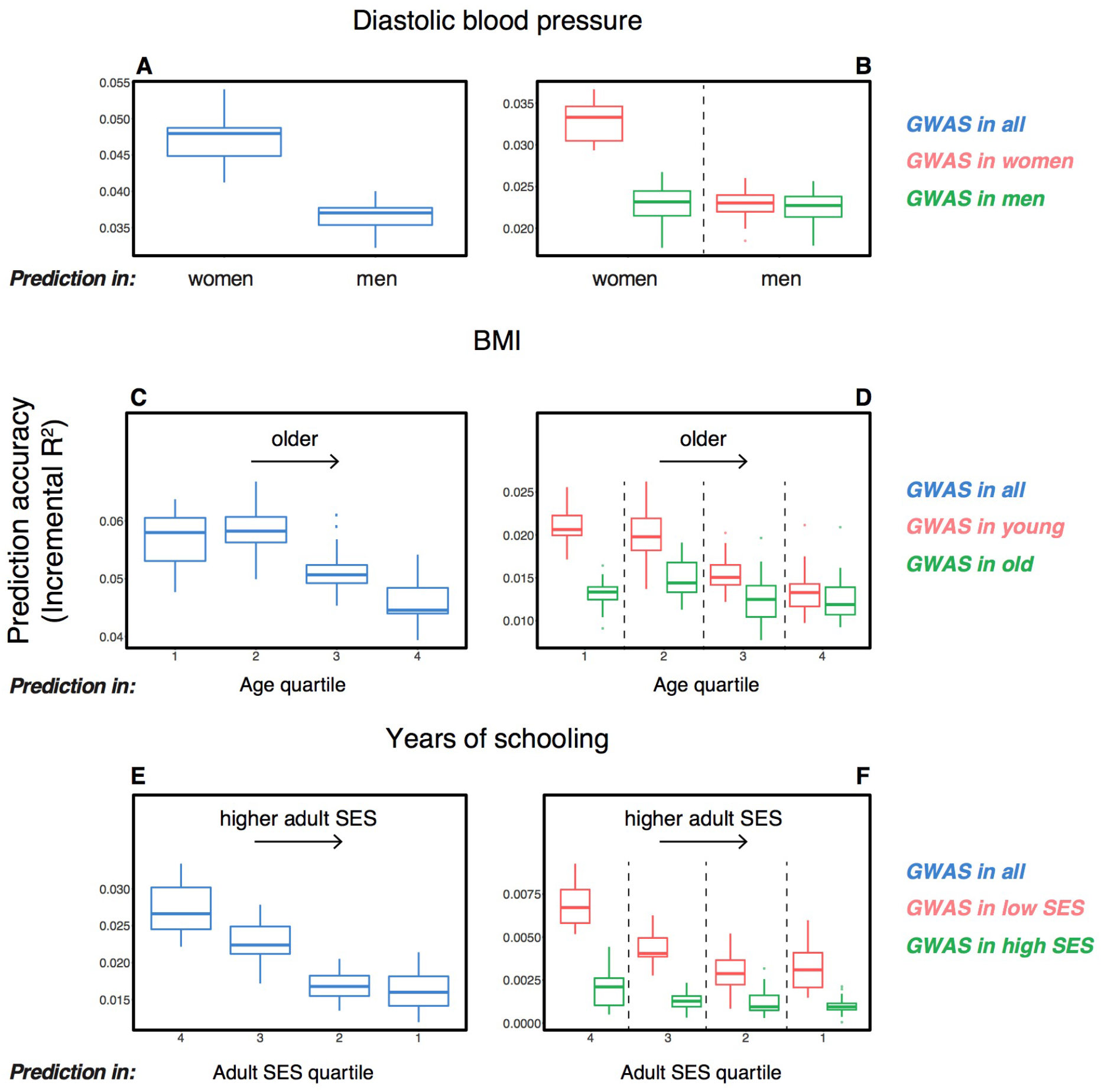
Variable prediction accuracy of polygenic scores within an ancestry group. Shown are incremental *R*^2^ values (i.e., the increment in *R*^2^ obtained by adding a polygenic score predictor to a model with covariates alone) in different prediction sets. Each box and whiskers plot is computed based on twenty choices of estimation and prediction sets. Thick horizontal lines denote the medians **(A,C,E)**. The polygenic scores were estimated in large samples of unrelated WB individuals. Phenotypes were then predicted in distinct samples of unrelated WB individuals, stratified by sex (A), age (C) or Townsend deprivation index, a measure of SES (E). **(B,D,F)** Same as in A,C,E, but here the polygenic scores are based on a GWAS in a sample limited to one sex, age or SES group. When the GWAS is performed in the group that showed higher prediction accuracy in A,C,E (women, young, low SES), the qualitative trend is the same; but when the GWAS is performed in men, old or high SES, prediction accuracy is diminished and similar across groups.

We then considered two other cases, evaluating prediction accuracy in groups stratified by age for BMI^1^ and by adult socio-economic status (SES) for years of schooling, using the Townsend deprivation index as a measure; our choices were motivated by prior evidence suggesting that these characteristics of the GWAS can influence heritability (Branigan, McCallum, and Freese 2013; Conley et al. 2015; Belsky et al. 2018; Ge et al. 2017; Elks et al. 2012). We withheld a random set of 10K individuals in each quartile of age and SES for prediction and performed GWAS using the remaining individuals, matching the sample sizes across quartiles in the GWAS set. Similar to our observation for diastolic blood pressure, the prediction accuracy varies across prediction sets: it is 1.25-fold higher for BMI in the youngest quartile compared to the oldest (Mann-Whitney *p* = 1.7 · 10^-8^; **Fig. 1C**), and 1.69-fold higher for years of schooling in the lowest SES quartile compared to the highest (Mann-Whitney *p* = 1.4 · 10^-11^; **Fig. 1E**). Furthermore, the differences across groups are again sensitive to the choice of the GWAS set: the differences are marked when GWAS is restricted to the youngest quartile for BMI and the lowest SES quartile for years of schooling, but diminished when the GWAS is performed in the oldest and the highest SES quartiles for BMI and years of schooling, respectively (**Figs. 1D,F**). These results remained qualitatively unchanged when we used *R*^2^ instead of incremental *R*^2^ to measure prediction accuracy (**Fig. S1**).

In these analyses, we used a p-value threshold of 10^-4^ for inclusion of a SNP in the PGS. The choice of how stringent to make the GWAS p-value threshold is important but somewhat arbitrary, with approaches ranging from requiring genome-wide significance to including all SNPs (Weedon et al. 2008; Pharoah et al. 2008; Euesden, Lewis, and O’reilly 2014; Vilhjálmsson et al. 2015; Ware et al. 2017; Mostafavi et al. 2017; Speidel et al. 2019). Often, this threshold is chosen to maximize prediction accuracy in an independent validation set. When the goal is to compare prediction performance across different groups, there is no obvious optimal choice of the p-value threshold^2^. As we show, however, the qualitative trends reported in **Fig. 1** do not depend on the p-value threshold choice (**Fig. S2**).

These results pertain to three exemplar traits and do not speak to the prevalence of this phenomenon. Nonetheless, they demonstrate that the portability of a polygenic score can vary markedly depending on sample characteristics of both the original GWAS and the prediction set, even within a single ancestry, and that the variation in prediction accuracy across strata can be substantial; in fact, on the same order as reported for different continental ancestries within the UK Biobank (Martin et al. 2019). As one example, the prediction accuracy in East Asian samples, averaged across a number of traits, is about half of that in European samples when GWAS was European-based; when the GWAS is done in the lowest SES group for years of schooling, prediction accuracy in the highest SES group is less than half of that in the lowest SES (**Fig. 1F**). Moreover, whereas for these traits, we had prior information about which characteristics may be relevant, other aspects that vary across sets of individuals are undoubtedly important as well (e.g., smoking behavior may modify genetic effects on lipid traits; Bentley et al. 2019), and for any given trait of interest, much less may be known *a priori.*

### Possible explanations for the variable prediction accuracy

Our goal in this paper is to highlight that prediction accuracies can vary across groups of highly similar ancestry, rather than to investigate the likely causes for any particular phenotype. Nonetheless, it is worth noting a couple of possibilities. Perhaps the simplest explanation for our findings is that prediction accuracies vary only because of differences in the extent of environmental variance, while the genetic variance is more or less constant. Indeed, the SNP heritabilities vary markedly across strata (see also Ge et al. 2017), and the prediction accuracies track heritability differences (**Fig. 2A,B,C**). For all three traits, however, the estimated SNP heritability increases or remains the same with increasing phenotypic variance, in contrast to what would be expected under a model with a fixed genetic variance across strata (**Fig. 2D,E,F**).

Another possibility is that there is an interaction between genetic effects and sample characteristics, for instance that different sets of genetic variants contribute to blood pressure levels in males and females or to BMI across different stages of life^3^. This explanation is not supported by bivariate LD-score regression, which indicates that the genetic correlations across strata are close to 1 (**Table S2; Materials and Methods**). Yet when we re-estimate individual SNP effects in the prediction sets for SNPs ascertained in the original GWAS, the estimated effects of trait-increasing alleles are larger in the groups with higher prediction accuracy (**Fig. S3; Materials and Methods**). A possible way to reconcile these findings is if effect sizes are highly correlated but systematically larger in the groups with higher prediction accuracy.

Other factors complicate interpretation, however, and may also contribute to our observations. In particular, for the case of educational attainment, conditioning on adult SES induces a form of range restriction, which could contribute to variable prediction accuracy across strata. We note, however, that we see highly variable prediction accuracies across SES strata even when the GWAS is conducted in all individuals (**Fig. 1E**); in that regard, our approach mimics what happens in practice when polygenic scores are used to predict phenotypes in a sample with a smaller range of SES (e.g., Rimfeld et al. 2018). More generally, although this type of range restriction is artificially amplified in our example, SES differences will often be a problem for GWAS in which the sample is not representative of the population; for instance, the most recent major GWAS of educational attainment (Lee et al. 2018) included numerous medical data sets and the 23andMe data set, which are not representative of the national population.

Another potentially important factor is that the adjustment for PCs may not be a sufficient control for the different ways in which population structure can confound GWAS results (Vilhjálmsson and Nordborg 2013), leading to variable prediction accuracy across strata if they differ in their population structure. To examine this possibility, we repeated the analysis in **Figs. 1B,D,F** but using a linear mixed model (LMM) approach (including PCs among other covariates; see **Materials and Methods**), and obtained qualitatively similar results (**Fig. S4**). Although not a perfect fix (Listgarten, Lippert, and Heckerman 2013; Mathieson and McVean 2013), the fact that we obtain similar results using PCs and LMM suggests that confounding due to population stratification in the UK Biobank alone does not explain the variable prediction accuracies across strata.

**Figure 2:**
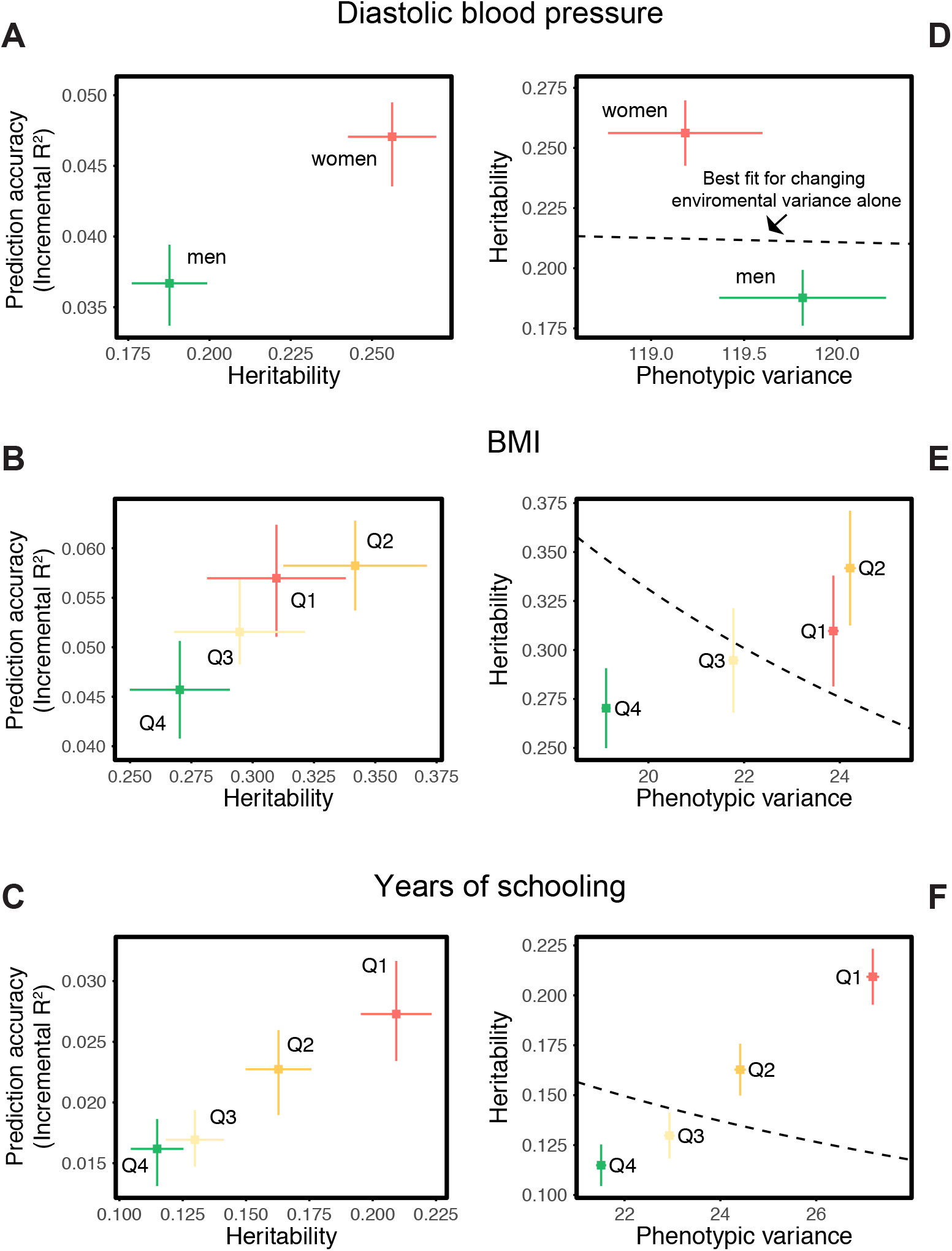
Differences in environmental variance alone do not explain the variable prediction accuracy. **(A,B,C)** The x-axes show heritability estimates (± SE) based on LD-score regression in each set. The y-axes show incremental *R*^2^ values as in Fig 1A,C,E. ‘Q’ denotes quartile of age and SES in (B) and (C), respectively. Throughout, prediction accuracy largely tracks SNP heritability. **(D,E,F)** The x-axes show phenotypic variance estimates (± SE) across strata after adjusting for covariates (sex, age and 20 PCs). If the heritability differences across strata are due to differences in environmental variance alone, with genetic variance constant, then heritability should be inversely proportional to phenotypic variance. However, the best fitting model for this inverse proportionality (dashed line) provides a poor fit.

### Potential portability obstacles explored through a comparison of standard and family-based GWAS

Beyond sample characteristics, a number of factors may shape the portability of scores across groups of similar ancestry. Standard GWAS is done in samples of individuals that deliberately exclude close relatives; as implemented, it detects direct effects of the genetic variants, but can also detect any indirect genetic effects of parents, siblings, or peers, effects of assortative mating among parents, and potentially environmental differences associated with fine scale population structure (Kong et al. 2018; Young et al. 2018; Lee et al. 2018; Trejo and Domingue 2019; Berg et al. 2019). Given that many of these effects are likely to be culturally mediated (e.g., Robinson et al. 2017; Ruby et al. 2018; Selzam et al. 2019), it seems plausible that they may vary within as well as across groups of individuals with different ancestries. To the extent that they contribute to GWAS estimates and hence to PGS, they may lead to differences in the prediction accuracy in samples unlike the original GWAS.

To demonstrate that these considerations are not just hypothetical, we compared the prediction accuracy when the PGS is trained on “unrelated” individuals such as those used in a standard GWAS to one obtained from a sibling-based (or “sib-based”) GWAS (**Materials and Methods**). In the latter, genotype differences between sibs—a result of random Mendelian segregation in the parents—are tested for association with the phenotypic difference between them. Because the tests depend on phenotypic differences between siblings who, of course, have the same parents, these tests are conditioned on the parental genotypes. Hence, they exclude many of the indirect effects signals that may be picked up in standard GWAS (**Supplementary Materials**). Differences between standard and sib-based GWAS are thus informative about the relative importance of factors other than direct genetic effects (Wood et al. 2014; Trejo and Domingue 2019; Lee et al. 2018; Berg et al. 2019; Selzam et al. 2019).

A challenge in this comparison is that the UKB contains about 22K sibling pairs, about 19K of which fall in the designation “White British” (WB). The siblings are similar to the unrelated individuals in terms of ages, SES distributions and genetic ancestries (**Figs. S5,S6**) but include a higher proportion of females; this difference is unlikely to influence our analyses (see below). While a large number, 19K pairs is still too few to have adequate power to discover trait-associated SNPs, when compared to a standard GWAS using the much larger sample of unrelated WB individuals (~340K).

To increase power and enable a direct comparison between the two designs, we split the SNP ascertainment and effect estimation steps as follows (**Fig. 3A**): we identified SNPs using a standard GWAS with a large sample size (median ~270K across the traits considered) (see **Materials and Methods**). We then estimated the effect of each significant SNP using (i) a sib-based association test and (ii) a standard association test. We chose the size of the estimation set in (ii) such that the median standard error of effect estimates in (i) and (ii) is approximately equal. We then compared the prediction accuracy of the two PGS obtained in this way (“standard PGS” and “sib-based PGS”) in an independent prediction set of unrelated individuals; as we show in the **Supplementary Materials**, our approach leads to highly similar prediction accuracies of the two approaches under a model with direct effects only (see **Materials and Methods** for details)^4^. A further advantage is that the two scores are compared for the same set of SNPs, such that LD patterns and allele frequencies do not come into play.

We applied the approach to 22 traits, focusing on traits with relatively high heritability estimates as well as social and behavioral traits that have been the focus of recent attention in social sciences. For the majority of the traits, such as diastolic blood pressure, BMI, and hair color, the prediction accuracies of standard and sib-based PGS were similar, as expected under standard GWAS assumptions and as observed for two traits simulated under these assumptions (**Fig. 3B**). However, for a range of social and behavioral traits, such as years of schooling completed, pack years of smoking and age at first sexual intercourse, the prediction accuracy of the sib-based PGS was substantially lower than that of the standard PGS (**Fig. 3B**). It was also significantly lower for two morphological traits, height and whole body water mass.

A number of factors could contribute to the difference between prediction accuracies for PGS based on sibs versus unrelated individuals, including residual effects of population stratification, indirect genetic effects from parents and assortative mating. The relative importance of each factor will vary across traits (Rosenberg et al. 2018; Kong et al. 2018; Haworth et al. 2019; Ruby et al. 2018; Selzam et al. 2019); for educational attainment, this gap is likely to reflect at least in part the documented contribution of indirect genetic effects to the standard PGS (Lee et al. 2018; Kong et al. 2018; Young et al. 2018). We show in the **Supplementary Materials** that in the presence of indirect genetic effects mediated through parents, standard PGS outperforms sib-based PGS unless direct and indirect effects are strongly anticorrelated (**Fig. S7**), which seems unlikely to be the case for years of schooling. The difference in the performance of sib-based and standard PGS observed for other social and behavioral outcomes, such as household income and age at first sexual intercourse (**Fig. 3B**), may reflect a similar phenomenon. An additional contribution to divergent prediction accuracies could come from sibling indirect effects, which contribute differentially to standard and sibling-based PGS.

For height, there may be an important contribution of assortative mating to the difference in prediction accuracies (Wood et al. 2014; Robinson et al. 2017; Lee et al. 2018). In the **Supplementary Materials**, we show that under a simple model of positive assortative mating (mating of similar individuals), the prediction accuracy based on a standard PGS is better than that of a sib-based PGS (**Fig. S8**). The difference in the performance of sib-based and standard PGS observed for whole body water mass (**Fig. 3B**) could possibly reflect the same underlying effects of assortative mating, especially considering the high genetic correlation between the two traits (by bivariate LD score regression, *ρ_g_* ≈ 0.66, *p* < 10^-30^). We further confirmed that the difference in the sex ratio of the siblings and unrelated individuals, mentioned earlier, has a negligible effect on these differences (**Fig. S9**).

**Figure 3:**
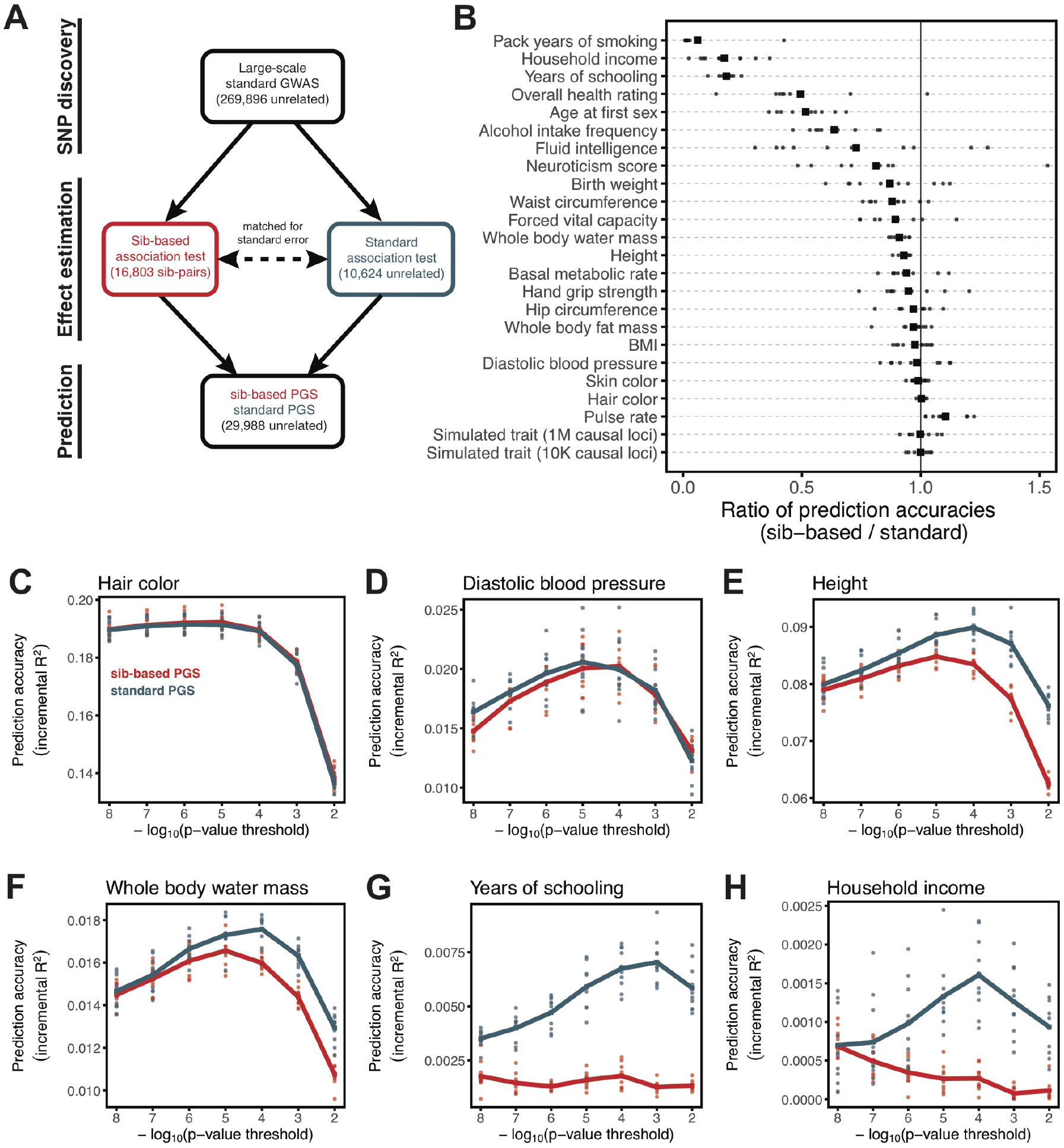
Comparison of prediction accuracy of standard and sib-based polygenic scores. **(A)** After ascertaining SNPs in a large sample of unrelated individuals, we estimated the effect of these SNPs with a standard regression using unrelated individuals and, independently, using sib-regression. We then used the polygenic scores for prediction in a third sample of unrelated individuals. We chose the sample size of the standard PGS estimation set such that median effect estimate SEs are equal in the two designs, thereby ensuring equal prediction accuracy under a vanilla model with no indirect effects or assortative mating. Numbers in parentheses are median sample size in each set across 22 traits in **Table S1** (see **Table S3** for sample sizes for each trait). **(B)** Ratio of prediction accuracy in the two designs across 22 traits. For each trait, we performed 10 resampling iterations of unrelated individuals into three sets for discovery, estimation and prediction (small points). Large points show mean values. **(C-H)** We repeated this procedure with different discovery-set p-value thresholds for including a SNP in the polygenic score. The higher the p-value threshold is, the more SNPs are included. For each p-value threshold, points show 10 iterations as described and lines show mean values. Shown are a subset of traits, with traits appearing in (B) but not shown here presented in **Fig. S10**.

Thus, in comparisons of the prediction accuracies for PGS derived from standard and sib-based association tests, many traits, notably behavioral ones, show substantial differences in performance. We caution that while lower prediction accuracies for PGS based on sib-based GWAS suggest that assortative mating or indirect effects play a substantial role, the magnitude of the ratio also depends on other features of the comparison like the sample sizes used (see **Supplementary Materials**). By matching the sampling errors of the two approaches (**Fig. 3A**), we ensure that prediction accuracies are comparable in the absence of complications such as assortative mating or indirect effects. But in the presence of these complications, the relative prediction accuracies will depend on sample sizes and on the contributions of environmental, direct and indirect genetic components to phenotypic variance. Indeed, we show in the **Supplementary Materials** that in the presence of indirect genetic effects or assortative mating, the difference in prediction accuracies between the two approaches stems in part from the noise-to-signal ratio for sib-based versus standard GWAS. An implication is that the gap between the prediction accuracy of sib-based and standard PGS should depend on the number of SNPs included in the polygenic scores (**Figs. S7,S8**).

Motivated by these considerations, we examined how the prediction accuracy varies when progressively relaxing the GWAS p-value threshold for inclusion of SNPs, i.e., when including more weakly associated SNPs in the PGS. (In **Fig. 3B**, results are shown for the p-value threshold that maximizes the prediction accuracy of the standard PGS, replicating the practice when comparing populations of different ancestry (Martin et al. 2019).) For hair color and blood pressure, there is little to no difference in prediction accuracy between the two estimation methods, regardless of the number of SNPs included in the score (**Figs. 3C,D**). In contrast, for height and whole body water mass, although standard and sib-based PGS perform similarly when based on the most significantly associated SNPs, standard PGS progressively outperforms sib-based PGS when more SNPs are included (**Figs. 3E,F**). Similarly, the difference in prediction accuracy between sib-based and standard PGS changes markedly for years of schooling, household income and other social and behavioral traits (**Figs. 3G,H and S10**). The growing gap in performance with increasing p-value threshold likely reflects a combination of an increasing noise-to-signal ratio in the sib-based PGS (see **Supplementary Materials**) and changes in the relative importance of direct effects versus other factors such as indirect parental effects and assortative mating.

In summary, the differences between the prediction accuracies of standard and sib-GWAS seen for a number of traits (**Fig. 3B**) demonstrate that standard GWAS estimates often include a substantial contribution of factors other than direct effects. In these cases, even if the power to detect direct effects were comparable, standard GWAS would lead to higher prediction accuracy than sib-GWAS. In some contexts that may be a sufficient reason to rely on PGS derived from standard GWAS. However, that gain stems from the inclusion of factors such as indirect effects and assortative mating that are likely to be modulated by SES, environment and culture (Selzam et al. 2019; Stulp et al. 2017). Thus, the increased prediction accuracy likely comes at a cost of not always porting well across groups, even of the same ancestry, in ways that may be difficult to anticipate.

## Implications

Although the conversation around the portability of PGS has largely focused on genetic ancestries, our results show that prediction accuracy can also differ, at times to a comparable extent, among groups of similar ancestry—even due to basic study design differences such as age and sex composition. If only due to increased environmental variance, such decreased accuracy would be acceptable, at least for certain applications. But as we have shown, differences in the degree of environmental variance are not the primary explanation for the patterns we report (**Fig. 2**), and other factors, including differences in the magnitude of genetic effects among groups, indirect effects and assortative mating, also lead to differences in the prediction accuracy of PGS, in ways that may make applications of phenotypic prediction problematic, even within a single ancestry group.

Following the discussion of portability across ancestries, we have focused on incremental R^2^ as a measure of portability, and it remains unknown to what extent the same issues also impact the use of PGS in reliably identifying individuals in the tails of the distribution, i.e., those at elevated risk of developing a disease—the main application of PGS in human genetics, as distinct from social science or evolutionary biology. Nonetheless, the same concerns are likely to apply, especially when the magnitude of genetic effects depends on GWAS characteristics.

In any case, these results make clear that the question of the domain over which a PGS applies is not just about population genetic parameters such as LD patterns and allele frequencies or GxG effects but also the extent of environmental variance, GxE, as well as the contribution of direct effects versus indirect effects, assortative mating and environmental confounding. An important implication is that differences in prediction accuracies among groups with distinct ancestries cannot be interpreted exclusively or even primarily in terms of population genetic parameters when these groups differ dramatically in their SES (Chetty et al. 2018; Conley 2010; Nuru-Jeter et al. 2018; Reich 2017) and other factors that may affect portability—especially when the relative contribution of these factors to GWAS signals remains unknown. Thus, efforts to conduct GWAS in groups that vary in ancestry and geographic locations will need to be accompanied by a careful examination of variation in portability along other dimensions.

In that regard, it is worth noting that while classical twin studies were often constituted to be representative of a reference population (often national in nature) (Branigan, McCallum, and Freese 2013; Polderman et al. 2015), the same is not true of most contemporary human genetic datasets, which are skewed towards medical case-control studies, biobanks that are opt-in (and thus tend to be wealthier and better educated than the population average) or direct-to-consumer proprietary genetic databases (which are even more skewed along these dimensions) (Lee et al. 2018). For instance, individuals in UK Biobank have higher SES than the rest of the British population (Fry et al. 2017) and are presumably self-selected for a certain level of interest in biomedical research. These factors alone raise challenges as to the broad portability of PGS derived from them.

One fruitful way forward may be to study data from related individuals, in which it should be possible to decompose the components of the signals identified in GWAS into direct and indirect effects, the degree of assortative mating and the contribution of residual stratification (Young et al. 2018; Kong et al. 2018; Zhang et al. 2015). Not only will this decomposition help us to better interpret the results of GWAS and the resulting PGS, it will make it possible to examine under which circumstances, and for which phenotypes, components port more reliably to other sets of individuals, both unrelated and related. Ultimately, we envisage that in order to be broadly applicable, GWAS-based phenotypic prediction models will need to include not only a PGS but some study characteristics, other social and environmental measures and, perhaps crucially, their interactions.

## Materials and Methods

### UK Biobank

The UK Biobank (UKB) is a large study of about half a million United Kingdom residents, recruited between 2006 to 2010 (Bycroft et al. 2017). In addition to genetic data, hundreds of phenotypes were collected through measurements and questionnaires at assessment centers, and by accessing medical records of the participants.

#### Inclusion criteria

In this study, we focused on 408,494 participants who passed quality control (QC) measures provided by UKB; specifically, for whom the reported sex (QC parameter “Submitted.Gender”) matched their inferred sex from genotype data (QC parameter “Inferred.Gender”); who were not identified as outliers based on heterozygosity and missing rate (QC parameter “het.missing.outliers”==0); and did not have an excessive number of relatives in the database (QC parameter “excess.relatives”==0). We further restricted ourselves to those individuals identified by UKB to be of “White British” (WB) ancestry (QC parameter “in.white.British.ancestry.subset”==1), which is a label that refers to those who, when given a set of choices, self-reported to be of “White” and “British” ethnic backgrounds and, in addition, were tightly clustered in a principal component analysis of the genotype data, as detailed in (Bycroft et al. 2017). For a given trait, we further conditioned on individuals for which measurement or report of the trait value was available.

### Phenotype data

We focused on 22 traits, including a range of well-studied physical, social, behavioral and health-related outcomes for which significant SNP heritabilities have been documented (see **Table S1** for a complete list of phenotypes, and their corresponding UK data field number). We calculated the phenotype “years of schooling” by converting the maximal educational qualification of the participants to years following Okbay et al. (Okbay et al. 2016) (**Table S4**). For diastolic blood pressure, pulse rate, and forced vital capacity, we took the average of the first two rounds of measurement taken during the same examination at UKB assessment centers. We adjusted the diastolic blood pressure levels for blood pressure lowering medication following Evangelou et al. (Evangelou et al. 2018) by shifting the values upward by 10 mm Hg for individuals taking medication. For hand grip strength, we took the average of the measurements for the two hands. The phenotype “household income” was defined as the average total household income before tax reported by the participants, categorized into five categories: less than £18,000, £18,000 to £29,999, £30,000 to £51,999, £52,000 to £100,000, and more than £100,000. For a subset of individuals, multiple measurements of a phenotype were provided, corresponding to multiple visits to UKB assessment centers; in those cases, we used the measurements during the first visit.

### Genotype data

UKB participants were genotyped on either of two similar genotyping arrays, UK Biobank Axiom and UK BiLEVE arrays, at a total of ~850K markers. We focused on autosomal bi-allelic SNPs shared between both arrays, and used *plink v. 1.90b5* (Chang et al. 2015) to filter SNPs with calling rate >0.95, minor allele frequency >10^-3^, and Hardy-Weinberg equilibrium test p-val>10^-10^ among the WB samples, resulting in 616,323 SNPs.

### GWAS and trait prediction methods

#### GWAS by sample characteristics

We focused on a set of 337,536 WB samples that were identified by the UKB to be “unrelated” (sample QC parameter “used.in.pca.calculation”==1 as provided by UKB), defined such that no pairs of individuals are inferred to be 3^rd^ degree relatives or closer. We split the sample into nonoverlapping sets of individuals by one of the following factors: age at recruitment (in years), sex, and Townsend deprivation index at recruitment (used as a proxy for socioeconomic status or SES). For the Townsend deprivation index and age, we divided into four sets: Q1 [minimum value, first quartile], group 2 (first quartile, second quartile], group 3 (second quartile, third quartile], and group 4 (third quartile, maximum value]. We randomly selected 10K samples in each SES and age group, and 20K of males and 20K of females as held-out prediction sets, and performed GWAS using the remaining samples, matching sample sizes across groups in the GWAS set. We performed nine GWAS: for years of schooling in SES Q1 and SES Q4 (sample size 73,298 for each), and in the pooled sample of all four groups (sample size 293,192); for body mass index (BMI) in Q11 and Q4 (sample size 72,343 for each), and in pooled sample of all four groups (sample size 272,508); and for diastolic blood pressure in males and females (sample size 122,791 for each), and in a pooled sample of males and females (sample size 245,582). We performed all GWAS using *plink v. 2.0* (with flag: --linear), adjusting for sex, age and first 20 PCs as covariates. PCs are principal components of all genotype data, not just WB, as provided by UKB. For a subset of cases, (where GWAS was performed in samples restricted by characteristics described above), we additionally performed association tests using a linear mixed model (LMM) as implemented in *BOLT-LMMv. 2.3.2* (Loh et al. 2015), using LD scores computed from 1000 Genomes European-ancestry samples, with sex, age and first 20 PCs as covariates. The GWAS summary statistics were used to construct PGS for the samples in the prediction sets.

To better understand the performance of PGS across the strata (see “**Possible explanations for the variable prediction accuracy**”), we estimated the mean effect sizes of significant SNPs in each strata. To avoid overfitting, we first performed an association test in the pooled sample of all strata; then for significantly associated SNPs, we re-estimated the effect sizes in each of the strata. We performed 20 iterations of all above steps (**Fig. 1, Fig. S1-S4**).

We also considered two binary phenotypes (i) attained a college degree or not and (ii) attained any degree or not, for the analysis of educational attainment by SES (as described above for years of schooling), confirming that our analysis is robust to how education phenotype is coded (**Fig. S11**). For these traits we used a logistic regression model for GWAS (using *plink v. 2.0* with flag: --logistic).

#### Standard versus sibling-based regression

We used the genetic relatedness information provided by UKB to infer sibling pairs among the WB samples. Following Bycroft et al. (2017), we marked pairs with 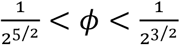 and IBS0 > 0.0012 as siblings, where *𝜙* is the estimated kinship coefficient and IBS0 is the fraction of loci at which individuals share no alleles. By this approach, we identified 19,335 sibling pairs including 35,464 individuals across 17,305 families. For a given trait, we included pairs with the property that trait values for both individuals were reported. We then formed two sets of individuals: “Siblings” set, including the sibling pairs randomly sampled to include only one pair per family, and an “Unrelateds” set, including the unrelated individuals identified by the UKB (see section GWAS by sample characteristics above), but excluding the Siblings and 7,409 individuals that were related to the Siblings (3rd degree or closer).

We focused on 22 traits (**Table S1**) and two simulated traits (see below). For each trait, we first downsampled the Unrelateds to a sample size *n*\* such that the median standard error of effect estimates roughly matched the median standard error in the sibling-based regression (see *“Estimating n*\*” below). We then divided the Unrelateds set into three non-overlapping sets: after sampling *n*\* individuals (Unrelateds-*n*\* set), we randomly split the rest of the Unrelateds set into an Unrelateds-prediction set (10% of the samples) to be used as a sample for trait prediction (“prediction set”), and an unrelated individuals discovery set (90% of the samples) to be used for the discovery of trait associated variants (see **Table S3** for sample sizes in each set). For each trait, we performed standard GWAS in the Unrelateds-discovery set, and ascertained SNPs by thresholding on association p-values. We then estimated the effect sizes for these ascertained SNPs in two ways: by a sibling-based association test in the Siblings set (using *plink v. 1.90b5’s* QFAM procedure; flag: --qfam), and by a standard association test in the Unrelateds-*n*\* set (using plink v. 2.0). Subsequently, for each set of ascertained SNPs in the Unrelateds-discovery set, two PGS were constructed for the samples in the Unrelateds-prediction set (see **Fig. 3A** for overview of the pipeline). We performed 10 iterations of the above sampling, ascertainment and estimation steps.

#### Estimating n*

In order to compare the performance of sibling-based and standard GWAS designs, we wanted to match both analyses to have similar prediction accuracy under a vanilla model of no assortative mating, population structure stratification or indirect effects. In the **Supplementary Materials**, we show that this could be achieved by matching median effect estimate standard errors. For each trait, we therefore calculated *n*\*, the sample size of a standard GWAS that yields roughly equal standard errors in the standard and sibling-based regressions. Specifically, for each trait, we first performed sibling-based GWAS in the Siblings using plink’s QFAM procedure (using the flag: -- qfam mperm=100000 emp-se). We then randomly sampled a range of sample sizes from the set of Unrelateds, from 5K to 20K in 1K increments. Following Wood et al. (Wood et al. 2014), for each sample size, we performed a standard GWAS, and investigated the linear relationship between the square root of the sample size and the inverse of the median standard error of the effect size estimates. We then used this linear relationship to estimate the sample size of a standard GWAS that corresponds to the inverse of the median standard error of the effect sizes estimate in the sibling-based GWAS.

All standard association tests were performed using *plink v. 2.0* (using the flag: --linear), adjusting for sex, age and first 20 PCs as covariates. For sibling-based association tests we first residualized the phenotypic values on the same covariates, and then regressed the sibling differences in residuals on sibling genotypic differences using plink’s QFAM procedure as described above.

We also considered a version of the analysis described above, in which we first residualized the phenotypes on covariates in the pooled sample of all WB individuals, and then ran the pipeline on the residuals without further adjustment for covariates in the GWAS or prediction evaluation. As shown in **Fig. S12**, this approach produced results that are qualitatively the same to what we present in **Fig. 3**.

#### Simulated traits

We wanted to check that given the study design described above, sibling-based and standard GWAS perform similarly with respect to trait prediction, under the vanilla model of no population stratification, assortative mating or indirect genetic effects (**Fig. 3**). To this end, we simulated two traits with (i) heritability *h*^2^ = 0.5 and *m* = 10,000 causal loci, and (ii) heritability *h*^2^ = 0.5 and *m* = 1,000,000 causal SNPs.

We randomly selected the causal SNPs from a set of 10,879,183 imputed SNPs, considering that most causal variants are plausibly not directly genotyped on SNP arrays. We used a set of SNPs that passed quality control procedures by the Neale lab (http://www.nealelab.is/uk-biobank), namely autosomal SNPs, imputed using the haplotype reference consortium (HRC) panel, which have INFO score > 0.8 and have minor allele frequency > 10^-4^; we further limited the SNP set to ones that were bi-allelic in the WB sample. As in Martin et al. (Martin et al. 2017), we randomly assigned effect sizes to these causal SNPs as 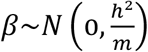, and zero for non-causal SNPs. We then calculated genetic component of the trait, *g*, for all WB samples under an additive model by summing the allelic counts weighted by their effect sizes using plink (using the flag: --score). Allelic counts were determined by converting imputation dosages to genotype calls with no hard calling threshold. We also assigned environmental contributions as *ε*~*N*(0,1 – *h*^2^), and then constructed the PGS for each individual,

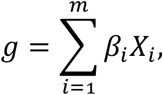

where *X_i_* is the number of minor alleles at SNP *i* carried by the individual, and the trait value for the individual is calculated as the sum of genetic and environmental contributions:

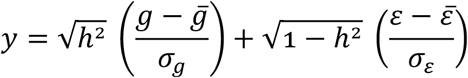

where bars represent averages, *σ_g_* is the standard deviation of PGS across individuals and *σ_ε_* is the standard deviation of environmental contributions across individuals. These simulated traits were then analyzed using the same pipelines as the other traits (e.g., adjusting for covariates etc.). Importantly, SNP discovery and effect size estimations in GWAS were performed without knowledge of the causal SNPs.

#### Polygenic score (PGS) construction and trait prediction

For all GWAS designs described above, we used p-value thresholding followed by clumping to choose sets of roughly independent SNPs to build PGS. We considered a logarithmically-spaced range of p-values: 10^-8^, 10^-7^, 10^-6^, 10^-5^, 10^-4^, 10^-3^, and 10^-2^ (or a subset if no SNP reached that significance level). We then used plink’s clumping procedure (using the flag: --clump) with LD threshold *r*^2^ < 0.1 (using 10,000 randomly selected unrelated WB samples as a reference for LD structure) and physical distance threshold of >1MB. The selected SNPs were then used to calculate PGS for individuals in the prediction sets, by summing the allelic counts weighted by their estimated effect sizes (log of the odds ratios in the case of binary traits) using plink (using the flag: --score). We calculated the incremental *R*^2^: we first determined *R*^2^ in a regression of the phenotype to the covariates, and then calculated the change in *R*^2^ when including the PGS as a predictor. For binary traits, we calculated incremental Nagelkerke’s *R*^2^.

#### Estimating heritability and genetic correlation

We calculated SNP heritability across sex, age and SES groups for diastolic blood pressures, BMI and years of schooling, respectively (as described in the section “GWAS by sample characteristics”) as well as genetic correlations across pairs of groups: we first performed GWAS using all unrelated WB individuals in each group. We then used the GWAS summary statistics to perform LD-score regression with LD scores computed from the 1000 Genomes European-ancestry samples (Bulik-Sullivan et al. 2015). We also calculated genetic correlation between height and whole body water mass, using all unrelated WB individuals for GWAS.

## Acknowledgements

We are grateful to Ipsita Agarwal, Daniel Belsky, Jeremy Berg, Graham Coop, Doc Edge, Iain Mathieson, Augustine Kong, Magnus Nordborg, Guy Sella, Alex Young and members of the Przeworski and Sella labs for valuable discussions and Ipsita Agarwal and Doc Edge for comments on a draft of the manuscript. This work was funded by NIH GM121372 to MP, NIH HG008140 to JKP and Robert Wood Johnson Foundation Pioneer Award (grant number 84337817) to DC.

## Footnotes

1 Since the UK Biobank participants were enrolled within about a five-year span, differences in age could in principle also be reflective of cohort effects.

2 The optimal p-value in this context will differ across studies, as it depends not only on the genetic architecture and heritability of the trait, but also on the GWAS sample size, i.e., power (Dudbridge 2013).

3 Although such interactions could in some contexts be thought of as reflecting GxE, we use the term sample characteristic rather than “environment”, as environment has different meaning across disciplines, referring in some contexts only to factors that are “exogenous” to genetics. Viewed in this lens, SES in adulthood cannot be interpreted as exogenous, because it is in part determined by educational achievement, which is itself influenced by genetic factors, and similarly it is questionable whether age or sex are environments.

4 Because the first step of our study design is to identify SNPs that are associated with the trait in a large set of unrelated individuals and we subsequently match the sampling variances of sib and standard GWAS, rather than identify distinct sets of SNPs separately in the two designs, the ratio of prediction accuracies that we obtain cannot be directly compared to those reported in other studies.

