## Supplementary Materials for "Variable prediction accuracy of polygenic scores within an ancestry group"

### Supplementary Materials for: Variable portability of polygenic scores within an ancestry group

May 7, 2019

### 1 Prediction accuracies of polygenic scores based on standard and sib-GWAS

#### 1.1 Overview of derived results

In the main text, we compare the prediction accuracies of polygenic scores (PGS) based on a standard GWAS of unrelated individuals and a GWAS based on sibling differences, for a number of traits. Here, we describe how this comparison is implemented, and how indirect effects and assortative mating manifest in this comparison.

**Matching standard and sib-based prediction accuracies.** Current standard GWAS are based on huge sample sizes, leading to less noisy estimates than are afforded by family association studies such as those based on sib differences, which are typically much smaller. This difference in precision needs to be taken into account in making comparisons between the prediction accuracy of scores derived from the two approaches. We show that under a vanilla additive model with no assortative mating, indirect effects, population structure (or other complications), and if the standard GWAS is subsampled to a sample size

$$n^* \approx \frac{1}{1+(1-h^2)(1-2\rho_{sibs})} n^{pairs},$$

where $n^{pairs}$ is the number of sib pairs, $h^2$ is the heritability and $\rho_{sibs}$ is the correlation in environmental effects experienced by siblings, the two study designs are expected to have the same (out-of-sample) prediction accuracy (see **Section 1.2**). This analytic result is not that useful in practice, however; in particular, it requires prior knowledge about the extent to which environmental effects correlate among siblings. Instead, we took an empirical approach to match the prediction accuracies in the two approaches: following [2], we subsampled the regular GWAS to match the median standard errors of the sib-GWAS. As we show in **Section 1.2.3**, under our vanilla model, we then expect near identical out-of-sample prediction accuracies for polygenic scores derived from the two study designs.

**Indirect parental effects.** In the presence of indirect parental effects, the out-of-sample prediction accuracy takes a simple form. For a polygenic score based on a standard GWAS, we obtain

$$E[R_{ur}^2] = \tau^2 \frac{1}{1+c},$$

where $\tau^2$ is the ratio of the variance in the trait due to both direct effects and indirect effects of transmitted parental alleles over the total phenotypic variance; and $c$ is a term representing the noise to signal ratio in a standard GWAS. For the polygenic score based on sib-GWAS, we obtain

$$E[R_{sib}^2] = (1 + \rho \frac{\sigma_\eta}{\sigma_\beta})^2 h_\beta^2 \frac{1}{1 + c\tau^2 / h_\beta^2}.$$

where $\sigma_\beta^2$ and $\sigma_\eta^2$ are the variances of random direct and indirect effects, respectively, $\rho$ is the correlation between direct and indirect effects, and $h_\beta^2$ is the proportion of the phenotypic variance explained by direct effects. Our results suggest that under plausible conditions, the presence of indirect effects would lead to higher prediction accuracy in a standard GWAS (after our sampling variance matching procedure). This result holds whether direct and indirect effects are positively correlated, uncorrelated or even somewhat negatively correlated (**Fig. S7**).

**Assortative mating.** We investigated several models of assortative mating by simulation. Standard GWAS-based polygenic scores have greater prediction accuracies than those based on sib-GWAS when the parental phenotypes are positively correlated, and the reverse is true if they are negatively correlated (**Fig. S8A,B**). The relative difference in prediction accuracies of the two study designs grows with the inclusion of more SNPs in the polygenic score model (**Fig. S8D,F**).

In our analytic considerations, we ignored the ascertainment step of our study design, in which it is decided which SNPs to include in the polygenic score. We assumed that SNPs are pre-ascertained and that the set of ascertained SNPs includes all causal ones. In a subset of simulations, we implemented the ascertainment step based on an independent simulated GWAS (see below). In both settings, we refer (somewhat loosely therefore) to the regression on ascertained SNPs in a sample of unrelated individuals as “standard GWAS” and the regression of the difference in phenotypes on the difference in sib genotypes as “sib-GWAS.”

#### 1.2 Picking the sample size of the standard GWAS sample to match the prediction accuracy of the score based on the sib-GWAS

We look for the sample size $n^*$ of a standard GWAS performed on sample of unrelated individuals such that, under our vanilla model, the resulting polygenic score has the same (out-of-sample) prediction accuracy as the polygenic score obtained from a sib-GWAS with sample size $n_{pairs}$. We begin by assuming that all causal sites $i$ are known; that they are unlinked; that they have only additive, direct effects on the phenotype; and that there is no population stratification or assortative mating. We first find the sampling variance of the effect size estimate for a single site obtained from each of the two study designs. We will then examine (and ultimately match) the prediction accuracy of the polygenic scores obtained from effect sizes estimated in the estimation sets, $\hat{\beta}_{ur}, \hat{\beta}_{sib}$, on a new, independent prediction sample of unrelated individuals $\{(x', y')\}$.

##### 1.2.1 Sampling error of the estimated effect size at a single site

Our model for the phenotypic value $y$ is

$$y = g + e$$

where $e$ is a Normally distributed environmental effect (which includes all sources of random noise) and

$$g = \beta_0^{ur} + \sum_i \beta_i x_i$$

where $x_i \in \{0, 1, 2\}$ are random genotypes. The genotype is coded as the the number of alleles with effect $\beta_i$ carried by the individual at site $i$. Effect sizes $\beta = \{\beta_i\}$ are treated as fixed parameters throughout (except when noted otherwise in the very last step leading to eq. 22). We can rewrite our model to focus on the effect size at a single site $i$:

$$y = \beta_0 + \beta_i x_i + \epsilon_i, \tag{1}$$

where

$$\epsilon_i = g - \beta_i x_i + e,$$

with variance

$$Var[\epsilon_i] = Var[g - \beta_i x_i] + Var[e] = Var[y] - \beta_i^2 Var[x_i]$$

In an OLS regression, the standard error for the effect of an allele at site $i$ is

$$Var[\hat{\beta}_i^{ur}] = \frac{Var[\epsilon_i]}{(n-1)Var[x_i]} = \frac{Var[y] - \beta_i^2 Var[x_i]}{(n-1)Var[x_i]}, \quad (2)$$

where $n$ is the sample size. In sib-GWAS, our model for site $i$ is

$$\Delta y = \beta_0^s + \beta_i \Delta x_i + \Delta \epsilon_i,$$

with variance

$$Var[\Delta\epsilon_i] = Var[\Delta g - \beta_i \Delta x_i] + Var[\Delta e] =$$

$$Var[\Delta g] + \beta_i^2 Var[\Delta x_i] - 2\beta_i^2 Var[\Delta x_i] + Var[\Delta e].$$

Recall that for siblings (denoted with subscripts $A$ and $B$), we expect

$$Cov[x_{i,A}, x_{i,B}] = \frac{1}{2} Var[x_i],$$

$$Cov[g_A, g_B] = \frac{1}{2} Var[g].$$

Plugging these back in, we obtain

$$Var[\Delta\epsilon_i] = Var[g] - \beta_i^2 Var[x_i] + 2Var[e](1 - \rho_{sibs})$$

where $\rho_{sibs} = Cor[e_A, e_B]$ is the correlation in environmental effects between sibs. The variance of the estimated effect size in sib-GWAS is therefore

$$Var[\hat{\beta}_i^s] = \frac{Var[\Delta\epsilon_i]}{(n_{pairs} - 1)Var[\Delta x_i]} = \frac{Var[y] - \beta_i^2 Var[x_i] + Var[e](1 - 2\rho_{sibs})}{(n_{pairs} - 1)Var[x_i]}. \quad (3)$$

##### 1.2.2 Sample size required for matched prediction accuracy

We measure prediction accuracy as the expected squared correlation between polygenic scores $\hat{g}$ and phenotypic values in an independent prediction set of unrelated individuals, denoted $\{(x', y')\}$,

$$E[R^2] = \frac{Cov^2[\hat{g}(x'), y']}{Var[y']Var[\hat{g}(x')]},$$

by requiring

$$E[R^2(standard\, GWAS\, with\, sample\, size\, n^*)] \stackrel{!}{=} E[R^2(sib\, regression\, with\, sample\, size\, n_{pairs})]$$

or equivalently

$$\frac{Cov^2[\hat{g}_{sib}(x'), y']}{Var[\hat{g}_{sib}(x')]} \stackrel{!}{=} \frac{Cov^2[\hat{g}_{ur}(x'), y']}{Var[\hat{g}_{ur}(x')]}. \tag{4}$$

We solve eq. 4 for a sample size $n^*$ to be used for estimation of the polygenic score in a standard GWAS that satisfies eq. 4. We note that if the vector of estimates $\hat{\beta}$ is given, then

$$Cov_{\{(x',y')\}}[y', \hat{g}(x')|\hat{\beta}] = Cov_{\{(x',y')\}}[g(x'), g(x') + \sum_i^m x'_i(\hat{\beta}_i - \beta_i)|\hat{\beta}] =$$

$$Var_{\{(x',y')\}}[g(x')|\hat{\beta}] + \sum_i^m Cov_{\{(x',y')\}}[\beta_i x'_i, (\hat{\beta}_i - \beta_i)x'_i|\hat{\beta}] = \sum_i^m Var[x'_i]\beta_i\hat{\beta}_i, \tag{5}$$

To incorporate randomness both in the estimation set (summarized by the Multivariate Normal distribution of $\hat{\beta}$) and the prediction set $\{(x', y')\}$, we apply the law of total expectation on eq. 4,

$$\frac{E_{\hat{\beta}}[Cov_{\{(x',y')\}}[\hat{g}_{sib}(x'), y']]}{\sqrt{E_{\hat{\beta}}[Var_{\{(x',y')\}}[\hat{g}_{sib}(x')]]}} \stackrel{!}{=} \frac{E_{\hat{\beta}}[Cov_{\{(x',y')\}}[\hat{g}_{ur}(x'), y']]}{\sqrt{E_{\hat{\beta}}[Var_{\{(x',y')\}}[\hat{g}_{ur}(x')]]}}. \tag{6}$$

Since for every $i$, we have

$$E[\hat{\beta}_i^{ur}] = E[\hat{\beta}_i^{s}] = \beta_i,$$

we obtain

$$E_{\hat{\beta}^s}[Cov[y', \hat{g}^s(x')|\hat{\beta}^s]] = \sum_i^m Var[x'_i]\beta_i^2 = E_{\hat{\beta}^{ur}}[Cov[y', \hat{g}^{ur}(x')|\hat{\beta}^{ur}]],$$

which turns the requirement of eq. 6 into

$$E_{\hat{\beta}^s}[Var_{\{(x',y')\}}[\hat{g}_{sib}(x')]] \stackrel{!}{=} E_{\hat{\beta}^{ur}}[Var_{\{(x',y')\}}[\hat{g}_{ur}(x')]],$$

or simply

$$\sum_i^m Var[X_i]Var[\hat{\beta}_i^{ur}] \stackrel{!}{=} \sum_i^m Var[X_i]Var[\hat{\beta}_i^{s}]. \tag{7}$$

Plugging the sampling variance results from eq. 2 and eq. 3 into eq. 7 and reordering, we obtain

$$\frac{n^* - 1}{n_{pairs} - 1} = \frac{\sum_i^m Var[y] - \beta_i^2 Var[x_i]}{\sum_i^m Var[y] - \beta_i^2 Var[x_i] + Var[e](1 - 2\rho_{sibs})},$$

or, assuming that the trait is polygenic such that $m >> 1$,

$$\frac{n^*}{n^{pairs}} \approx \frac{1}{1 + (1 - h^2)(1 - 2\rho_{sibs})}. \tag{8}$$

Eq. 8 can in principle be applied to the estimation of $\rho_{sibs}$ for a given trait, under our model assumptions, and given an independent estimate of $h^2$.

##### 1.2.3 Empirical matching of standard errors

The result of eq. 8 is the same as we would obtain if we required

$$\forall i \; Var[\hat{\beta}_i^{sib}(x_i)] \stackrel{!}{=} Var[\hat{\beta}_i^{ur}(x_i^{sib})] \tag{9}$$

without taking into account randomness in the prediction set. In practice (and in the results shown in the main text), we have no prior knowledge about $\rho_{sibs}$ and instead we find a sample size $n^*$ for the standard GWAS such that

$$median_i(Var[\hat{\beta}_i^{sib}(x)]) \stackrel{!}{=} median_i(Var[\hat{\beta}_i^{ur}(x)]) \tag{10}$$

We note that the condition in eq. 9 is approximately met because, if we assume that $y$ is a highly polygenic trait where

$$\forall i \; \beta_i^2 Var[xi] << Var[y],$$

then

$$\forall i \; Var[\hat{\beta}_i^{sib}(x)] = Var[\hat{\beta}_i^{ur}(x)] = \frac{D}{Var[x_i]}$$

where $D$ is the same for sib-GWAS and standard GWAS estimates (given that a sample of $n^*$ is used), and approximately independent of $\beta_i$. Eq. 10 can be thought of as a weighted-median of

$D$. In conclusion, the requirement of eq. 10 leads to equal prediction accuracy of standard and sib-GWAS under the vanilla model assumptions. We note further that in the main text (**Fig. 3**), to follow common practice, we use incremental $R^2$ throughout rather than $R^2$; However, as we show in **Fig. S12**, the use of $R^2$ instead gives highly similar qualitative results.

#### 1.3 Indirect parental effects

##### 1.3.1 Distribution of the effect size estimate at a single site

We consider an additive model with direct effects as well as indirect parental effects, assuming no interaction between the parents and the polygenic score of the children and ignoring possible indirect effects of siblings on each other. We start by considering the model

$$y = \beta_0 + g + n + e$$

where $g$ is the sum of direct effects in an individual with genotype (effect-allele count) $x_i$ at each site $i$,

$$g = \sum_i^m \beta_i x_i,$$

and

$$n = \sum_i^m \eta_i (x_i + \tilde{x}_i^m + \tilde{x}_i^p)$$

is the sum of parental indirect effects, with parental allele counts $x_i + \tilde{x}_i^p + \tilde{x}_i^m$ at each site, where $\tilde{x}_i^m$ is the untransmitted maternal effect allele count, and $\tilde{x}_i^p$ the untransmitted paternal effect allele count, with $\tilde{x}_i^m, \tilde{x}_i^p \in \{0, 1\}$. As we show, when we choose the standard GWAS sample size $n^*$ such that the effect size estimate sampling errors match those of the sib-GWAS, the prediction accuracies of the two polygenic scores differ in an independent sample: unless there is a large negative correlation between indirect and direct effects, the polygenic score from standard GWAS is expected to outperform the one based on sib-GWAS.

We first examine the distribution of an estimated effect size of $x_i$ on the phenotype. The OLS regression for a single site in a standard GWAS follows eq. 1 and can be rewritten as

$$y = \beta_0 + (\beta_i + \eta_i) x_i + \eta_i (\tilde{x}_i^p + \tilde{x}_i^m) + \epsilon_i \tag{11}$$

with

$$\epsilon_i = g + n + e - (\beta_i + \eta_i) x_i - \eta_i (\tilde{x}_i^p + \tilde{x}_i^m).$$

By the assumption of no assortative mating or other population structure,

$$Cov[\tilde{x}_i^p, \tilde{x}_i^m] = Cov[x_i, \tilde{x}_i^m] = Cov[x_i, \tilde{x}_i^p] = 0. \tag{12}$$

It directly follows that under the generative model specified by eq. 11, the OLS regression of $y$ to $x_i$ and $\tilde{x}_i^p + \tilde{x}_i^m$ is a regression involving two independent variables. Therefore, $\hat{\beta}_i^{ur}$ is Normally distributed with expectation

$$E[\hat{\beta}_i^{ur}] = \beta_i + \eta_i.$$

We next calculate the variance of $\hat{\beta}_i^{ur}$. From assumption 12 and

$$Var[\tilde{x}_i^m + \tilde{x}_i^p] = Var[x_i],$$

we obtain

$$Var[\epsilon_i] = Var[y] + (\beta_i + \eta_i)^2 Var[x_i] + \eta_i^2 Var[x_i] - 2Cov[g+n, (\beta_i+\eta_i)x_i] - 2Cov[n, \eta_i(\tilde{x}_i^m + \tilde{x}_i^p)] =$$

$$= Var[y] - Var[x_i](\beta_i^2 + 2\beta_i\eta_i + 2\eta_i^2).$$

Finally,

$$Var[\hat{\beta}_i^{ur}] = \frac{Var[\epsilon_i]}{(n-1)Var[x_i]} = \frac{Var[y] - Var[x_i](\beta_i^2 + 2\beta_i\eta_i + 2\eta_i^2)}{(n-1)Var[x_i]}.$$

In sib regression, we have

$$\Delta y = \Delta g + \Delta e$$

since indirect parental effects cancel out when taking the difference between sibs (as siblings have an equal parental effect allele count). Thus, the expected estimate is the same as it was in the absence of indirect effects. Using the same considerations as in **Section 1.2** for the variance in sib differences, we obtain

$$\hat{\beta}_i^{sib} \sim N(\beta_i, \frac{Var[g] - \beta_i^2 Var[x_i] + Var[e](1 - 2\rho_{sibs})}{(n_{pairs} - 1)Var[x_i]}),$$

where $\rho_{sibs}$ is again the correlation in environmental effects between siblings.

##### 1.3.2 Polygenic score prediction accuracy

We now examine the difference in prediction accuracy of $\hat{g}^{ur}$ and $\hat{g}^{sib}$ after matching

$$Var[\hat{\beta}_i^{ur}] \stackrel{!}{=} Var[\hat{\beta}_i^{sib}] \tag{13}$$

by choosing a standard GWAS sample size $n^*$ that empirically satisfies the condition, as we do in the main text (see also **Section 1.2.3**). We can derive the expected prediction accuracy by averaging over both the estimation set (which we again shorthand as the distribution of $\hat{\beta}$) and the prediction set $\{(x', y')\}$. By the law of total expectation,

$$E[R^2] = E_{\hat{\beta}}[E_{\{(x',y')\}}[R^2]] \approx \frac{E_{\hat{\beta}}[Cov_{\{(x',y')\}}[\hat{g}(x'), y'|\hat{\beta}]]}{E_{\hat{\beta}}[Var_{\{(x',y')\}}[y']]E_{\hat{\beta}}[Var_{\{(x',y')\}}[\hat{g}(x')|\hat{\beta}]]}, \tag{14}$$

where we approximate the expected prediction accuracy by its first-order Taylor expansion. The numerator of eq. 14 is

$$E_{\hat{\beta}}[Cov_{\{(x',y')\}}[\hat{g}(x'), y'|\hat{\beta}]] = \sum_i^m (\beta_i + \eta_i) E_{\hat{\beta}}[Cov_{\{(x',y')\}}[\hat{\beta}_i x'_i, x'_i]|\hat{\beta}] = \sum_i^m Var[x_i](\beta_i + \eta_i)E[\hat{\beta}_i], \tag{15}$$

and the denominator terms are

$$E_{\hat{\beta}}[Var_{\{(x',y')\}}[y'|\hat{\beta}]] = Var[y] \tag{16}$$

and

$$E_{\hat{\beta}}[Var_{\{(x',y')\}}[\hat{g}(x')|\hat{\beta}] = E_{\hat{\beta}}[\sum_i^m Var[x_i]\hat{\beta}_i^2] = \sum_i^m Var[x_i](E[\hat{\beta}_i]^2 + Var[\hat{\beta}_i]). \tag{17}$$

Plugging eq. 15,16,17 back into eq. 14, we obtain

$$E[R^2] \approx \frac{(\sum_i^m Var[x_i](\beta_i + \eta_i)E[\hat{\beta}_i])^2}{Var[y](\sum_i^m Var[x_i]Var[\hat{\beta}_i] + \sum_i^m Var[x_i]E[\hat{\beta}_i]^2)}. \tag{18}$$

We note that

$$\tilde{C} := Var[y] \sum_i^m Var[x_i]Var[\hat{\beta}_i]$$

is the same for sib-GWAS and standard GWAS under the requirement of eq. 13. We therefore have

$$E[R^2_{ur}] \approx \frac{(\sum_i^m Var[x_i](\beta_i + \eta_i)^2)^2}{\tilde{C} + Var[y]\sum_i^m Var[x_i](\beta_i + \eta_i)^2}, \tag{19}$$

and

$$E[R^2_{sib}] \approx \frac{(\sum_i^m Var[x_i](\beta_i + \eta_i)\beta_i)^2}{\tilde{C} + Var[y]\sum_i^m Var[x_i]\beta_i^2}. \tag{20}$$

If we denote the proportion of the phenotypic variance explained by direct effects by

$$h^2_\beta := \frac{\sum_i^m Var[x_i]\beta_i^2}{Var[y]},$$

the proportion of the phenotypic variance explained by indirect effects of transmitted parental alleles by

$$\tau^2_\eta := \frac{\sum_i^m Var[x_i]\eta_i^2}{Var[y]},$$

and the proportion of phenotypic variance explained by both direct and indirect effects of transmitted alleles by

$$\tau^2 := \frac{\sum_i^m Var[x_i](\beta_i + \eta_i)^2}{Var[y]}$$

then the square of eq. 19 can be written as

$$E[R^2_{ur}] \approx \tau^2 \frac{1}{1+c}, \tag{21}$$

where we defined

$$c := \frac{\sum_i^m Var[x_i]Var[\hat{\beta}_i]}{\sum_i^m Var[x_i](\beta_i + \eta_i)^2}.$$

Here, $c$ can be thought of as a summary of the noise-to-signal ratio—with respect to the signal coming from both direct and indirect effects of transmitted alleles. If we treat effects $\beta$ and $\eta$ as random, treating results obtained thus far as conditional on $\beta$ and $\eta$, and further assume that effects are i.i.d. across sites (implying, in particular, that effect sizes and allele frequencies are independent),

$$\begin{pmatrix} \beta_i \\ \eta_i \end{pmatrix} \sim \left( \begin{pmatrix} 0 \\ 0 \end{pmatrix}, \begin{pmatrix} \sigma^2_\beta & \rho\sigma_\beta\sigma_\eta \\ \rho\sigma_\beta\sigma_\eta & \sigma^2_\eta \end{pmatrix} \right),$$

the expectation of the numerator of eq. 20 is

$$E_{\beta,\eta}[\sum_i^m Var[x_i]\beta_i(\beta_i+\eta_i)|\beta,\eta] = \sum_i^m Var[x_i]E_{\beta_i,\eta_i}[\beta_i^2+\beta_i\eta_i] = \sum_i^m Var[x_i](\sigma_\beta^2+\rho\sigma_\beta\sigma_\eta)$$

and thus eq. 20, in expectation, is:

$$E[R_{sib}^2] \approx E_{\beta,\eta}[E[R_{sib}^2|\beta,\eta]] = (1+\rho\frac{\sigma_\eta}{\sigma_\beta})^2 h_\beta^2 \frac{1}{1+c/\alpha}. \tag{22}$$

where

$$\alpha := h_\beta^2/\tau^2 = \frac{\sum_i^m Var[x_i]\beta_i^2}{\sum_i^m Var[x_i](\beta_i+\eta_i)^2}.$$

We examined the fit of this prediction to simulated data. Specifically, we ran simulations to estimate effect sizes in a sib-GWAS and in a standard GWAS, after choosing $n^*$ to match their sampling variances. Finally, we used the polygenic scores to predict phenotypic values in a sample of unrelated individuals (see **Section 1.3.3** for further detail).

**Fig. S7A,C,D** show the analytic result alongside simulation results, for different correlation coefficients between indirect and direct effect sizes. Even in the absence of a correlation between indirect and direct effect sizes, the polygenic score based on standard GWAS outperforms the polygenic score based on sib-GWAS.

To understand this behavior and dependency of the $\frac{R_{sib}^2}{R_{ur}^2}$ ratio on other parameters, we divide eq. 22 by eq. 21 and obtain

$$E[\frac{R_{sib}^2}{R_{ur}^2}] \approx \left(1+\rho\frac{\sigma_\eta}{\sigma_\beta}\right)^2 \alpha\frac{1+c}{1+c/\alpha}.$$

Noting further that

$$\left(1+\rho\frac{\sigma_\eta}{\sigma_\beta}\right)^2\alpha = \left(\frac{\sigma_\beta+\rho\sigma_\eta}{\sigma_\beta}\right)^2 \frac{\sigma_\beta^2}{\sigma_\beta^2+2\rho\sigma_\beta\sigma_\eta+\sigma_\eta^2} = 1-(1-\rho^2)\frac{\tau_\eta^2}{\tau^2},$$

we obtain

$$E[\frac{R_{sib}^2}{R_{ur}^2}] \approx [1-(1-\rho^2)\frac{\tau_\eta^2}{\tau^2}]\frac{1+c}{1+c\frac{\tau^2}{h_\beta^2}}. \tag{23}$$

A few conclusions emerge from eq. 23 and from the accompanying simulations. First, the sib-GWAS based polygenic score will outperform the standard GWAS-based polygenic score only if direct and indirect effects are strongly negatively correlated (see **Fig. S7A-D** for illustration). Second, the term

$$\frac{1+c}{1+c\frac{\tau^2}{h_\beta^2}} = \frac{1+\frac{\sum_i^m Var[\hat{\beta}_i]Var[x_i]}{\tau^2}}{1+\frac{\sum_i^m Var[\hat{\beta}_i]Var[x_i]}{h_\beta^2}} \tag{24}$$

can be interpreted as the dependence on the noise-to-signal ratio (where the signals are the proportions of phenotypic variance explained by direct and indirect effects of transmitted alleles). For a given sampling variance (matched across the two study designs), the extent of the signal will differ between standard GWAS and sib-GWAS. Importantly, the sampling variance influences the ratio of prediction accuracies. If indirect effects do not exist or make negligible contributions to the trait in question, then the ratio of prediction accuracies is expected to be close to one. In the presence of indirect effects, however, the magnitude of the deviation from 1 depends on the relationship between direct and indirect effects (and their covariance) as well as on the (matched) sampling variance. simulations of several parameter combinations suggest that the overall effect of this dependence on the noise-to-signal ratio is a decrease in $R^2_{sibs}/R^2_{ur}$ as noise increases; as more SNPs are included in the polygenic scores, the advantage of the standard GWAS-based polygenic score over the sib-GWAS based one grows larger (**Fig. S7E-H**). These considerations provide a possible interpretation for the patterns observed in **Fig. 3C-H** of the main text.

##### 1.3.3 Simulations of indirect effects

For each set of simulated individuals (discovery, estimation and prediction sets), we first simulated mother-father pairs, assigning parental alleles as $Bernoulli(p_i)$, where $p_i$ denotes the effect allele frequency at site $i$. We then sampled the parental alleles at random to generate offspring (one offspring per each mother-father pair to simulate a sample of unrelated individuals and two offspring to generate sibling pairs). Phenotypes of the offspring were assigned under an additive model, sampling from a Normal distribution with mean

$$\sum_{i}^{m} \beta_i x_i + \eta_i (x_i^p + x_i^m)$$

(where $x_i^m$ and $x_i^p$ are the maternal and paternal effect allele counts, respectively) and variance $\sigma_e^2$, representing the total variance of environmental effects. When there is no correlation between direct and indirect effects, $\sigma_e^2 = 1 - h_\beta^2 - 2\tau_\eta^2$. Using this approach, we generated a set of sibling pairs and estimated SNP effect sizes from these simulated data using a sib-GWAS. We calculated $n^*$ as follows: we simulated sets of unrelated individuals with a range of sample sizes. In each set, we performed a simple linear regression of the phenotypic values on the genotypes. We then estimated a linear relationship between the inverse of the median standard error of effect size estimates (as a dependent variable) and the square root of the sample size. Using this linear relationship, we predicted the sample size for the unrelated set that gives a median standard error equal to the median standard error of sib-GWAS effect size estimates ($n^*$). Finally, we simulated a set of unrelated individuals with sample size $n^*$ and compared the prediction accuracy ($R^2$) of the polygenic score based on standard GWAS on this sample with the one obtained from sib-GWAS.

We additionally investigated the effect of the number of SNPs included in the polygenic scores. For this analysis, we sorted the SNPs based on the association p-value obtained in an independent simulated set of unrelated individuals.

In these simulations, we used the following parameter values:

- The ratio of the phenotypic variance accounted for by direct effects versus by indirect effects ($h_\beta^2/\tau_\eta^2$): 5

- The phenotypic variance explained by offspring and parental alleles, given no correlation between direct and indirect effects ($h_\beta^2 + 2\tau_\eta^2$): 0.25 or 0.5

- The ratio of the variance of direct effects to the variance of indirect effects ($\sigma_\beta^2/\sigma_\eta^2$): 5

- Allele frequencies are drawn from a truncated exponential distribution, truncated on the left such that the minimum allele frequency is $1\%$.

- The number of loci, assumed independent (i.e., in linkage equilibrium): 100 (all causal), or 10,000 (all causal) or 10,000 (20% causal)

- SNP effect sizes drawn as

$$\left(\begin{array}{c}\beta_i\\ \eta_i\end{array}\right) \sim N\left(\left(\begin{array}{c}0\\ 0\end{array}\right), \left(\begin{array}{cc}\sigma_\beta^2 & \rho\sigma_\beta\sigma_\eta\\ \rho\sigma_\beta\sigma_\eta & \sigma_\eta^2\end{array}\right)\right),$$

where $\rho$ is the correlation between direct and indirect effect sizes. Effects sizes were then re-scaled to satisfy $\sum_i^m 2\beta_i^2 p_i(1-p_i) = h_\beta^2$ and $\sum_i^m 2\eta_i^2 p_i(1-p_i) = \tau_\eta^2$. Effects were set to 0 for non-causal loci.

- The number of sibling pairs for sib GWAS: 10,000

- The number of unrelated individuals for prediction: 10,000

- The number of unrelated individuals for discovery GWAS (i.e., to decide which SNPs to include): 20,000

- Number of iterations used to estimate $n^*$ and $R^2$ for a given set of parameters: 10

#### 1.4 Assortative mating

We consider assortative mating with regard to a phenotype, whereby the parents of individuals were more likely to mate if they were similar with respect to that phenotype. This process generates a correlation between genetic variants that contribute to the phenotype (i.e., linkage disequilibrium). Consequently, in a standard GWAS, the effect sizes of causal SNPs will partially capture the effect of other causal SNPs as well. Estimated effect sizes are thus expected to be inflated under positive assortative mating (mating of similar individuals) and deflated under negative assortative mating (mating of dissimilar individuals). In turn, in a sib-GWAS, the estimates are in expectation unaffected by assortative mating, because genetic differences between siblings arise from random Mendelian segregation in the parents.

##### 1.4.1 Simulations of assortative mating

We used simulations to examine the phenotypic prediction accuracies of polygenic scores based on sib- and standard GWAS under a model with assortative mating (assuming no indirect effects or population stratification beyond assortative mating); this end, we considered a sample of unrelated individuals, varying the degree of correlation between parental phenotypes $\rho_a$. Similar to our simulations for indirect effects (**Section 1.3.3**), we first simulated the estimation procedure in a sibling-based and in a standard GWAS (with sample size $n^*$). We then computed the prediction accuracy $R^2$ in an independent sample of unrelated individuals (see **Further simulation details** below).

We first considered the simple case of a single generation of assortative mating. In the presence of positive assortative mating ($\rho_a > 0$), polygenic scores based on standard GWAS outperform those based on sib-GWAS, whereas the opposite is true in the case of negative assortative mating ($\rho_a < 0$) (**Fig. S8A**). In simulations of two generations of assortative mating, the gap between the prediction accuracies of scores based on standard and sib-GWAS (**Fig. S8B**) widens, suggesting that our qualitative findings apply to scenarios of sustained assortative mating as well.

We further investigated prediction accuracy as a function of the number of SNPs included in the polygenic scores, by progressively increasing the p-value threshold, using p-values obtained from an independent GWAS in unrelated samples (similar to our analysis in **Fig. 3**). We considered two genetic architectures scenarios: (i) in which all SNPs are causal and (ii) the case in which 20% of of SNPs are causal (leading polygenic scores to include non-causal SNPs). Under both scenarios, the gap in prediction accuracies between standard and sib-GWAS grows with the number of SNPs (**Fig. S8C-F**).

**Further simulation details.** We simulated parental and offspring alleles as described for indirect effects in **Section 1.3.3**. To mimic assortative mating between parents, we first simulated i.i.d. genotypes (with effect allele counts $x_i$ at each SNP $i$) and randomly assigned "mother" and "father" labels to each individual. We then generated corresponding parental phenotypes under an additive model as

$$N(\sum_{i}^{m} \beta_i x_i, \sqrt{1-h^2})$$

where $\beta_i$ is the effect size of SNP $i$, and $h^2$ is the heritability. The same model was used to generate offspring phenotypes.

To induce a given correlation between parental phenotypes, $\rho_a$ (mimicking the consequence of the assortative mating process), we paired mothers and fathers as follows: First, we generated a random matrix

$$\begin{pmatrix} u_{m,i} \\ u_{p,i} \end{pmatrix} \sim N\left(\begin{pmatrix} \overline{y}_m \\ \overline{y}_p \end{pmatrix}, \begin{pmatrix} \sigma_{y_m}^2 & \rho_a \sigma_{y_m} \sigma_{y_p} \\ \rho_a \sigma_{y_m} \sigma_{y_p} & \sigma_{y_p}^2 \end{pmatrix}\right),$$

where $\overline{y}_m$ and $\overline{y}_p$ are the average phenotypes of mothers and fathers, respectively, $\sigma_{y_m}$ and $\sigma_{y_p}$ are the standard deviation of the phenotypes of mothers and fathers, respectively. We then sorted the mothers and fathers sets such that the ranks of values in $y_m$ and $y_p$ match the ranks of values in $u_m$ and $u_p$, respectively, to obtain $cor(y_m, y_p) \approx cor(u_m, u_p) = \rho_a$. In the case of two generations of assortative mating, we simulated the generation of the grandparents similarly. We compared the performance of polygenic scores based on standard and sib-GWAS as described in **Section 1.3.3**. In the simulations, we used the following parameter values:

- Heritability under random mating: 0.5

- The number of loci, assumed independent (i.e., in linkage equilibrium) under random mating: 10,000 (all causal) or 10,000 (20% causal)

- Allele frequencies, $p$, drawn from a truncated exponential distribution, truncated on the left such that the minimum allele frequency is $1\%$.

- SNP effect sizes set to 0 for non-causal loci and drawn as $\beta_i \sim N(0, \sigma^2)$, choosing $\sigma^2$ to satisfy $\sum_i^m 2\beta_i^2 p_i(1-p_i) = h^2$ for causal loci.

- The number of sibling pairs for sib GWAS: 10,000

- The number of unrelated individuals for prediction: 10,000

- The number of unrelated individuals for discovery GWAS (i.e., to decide which SNPs to include in the polygenic score): 20,000

- The number of iterations used to estimate $n^*$ and $R^2$ for a given set of parameters: 10

Table S1: Traits analyzed and their corresponding data fields in UK Biobank. For diastolic blood pressure, in addition to the raw measurements, we used medication data to adjust the blood pressure levels. For hand grip strength, we used the measurements for both hands.

| **Trait** | **UKB data field** |
|---|---|
| Age at first sex | 2139 |
| Alcohol intake frequency | 1558 |
| Basal metabolic rate | 23105 |
| Birth weight | 20022 |
| Body mass index | 21001 |
| Diastolic blood pressure | 4079, 6153, 6177 |
| Fluid intelligence | 20016 |
| Forced vital capacity | 3062 |
| Hair color | 1747 |
| Hand grip strength | 46, 47 |
| Height | 50 |
| Hip circumference | 49 |
| Household income | 738 |
| Neuroticism score | 20127 |
| Overall health rating | 2178 |
| Pack years of smoking | 20161 |
| Pulse rate | 102 |
| Skin color | 1717 |
| Waist circumference | 48 |
| Whole body fat mass | 23100 |
| Whole body water mass | 23102 |
| Years of schooling | 6138 |

Table S2: Genetic correlations across samples that vary by a study characteristic. Numbers are genetic correlations estimated using LD score regression for BMI, years of schooling and diastolic blood pressure, across samples stratified by age, socioeconomic status (SES) and sex, respectively. 'Q' denotes quartile of age or SES.

| **Trait/characteristic** | **Pair of strata** | **Genetic correlation (s.e.)** |
|---|---|---|
| BMI/Age | (Q1,Q2) | 0.93 (0.036) |
| | (Q1,Q3) | 0.95 (0.035) |
| | (Q1,Q4) | 0.95 (0.039) |
| | (Q2,Q3) | 0.89 (0.032) |
| | (Q2,Q4) | 0.91 (0.036) |
| | (Q3,Q4) | 1.00 (0.040) |
| Years of schooling/SES | (Q1,Q2) | 0.98 (0.054) |
| | (Q1,Q3) | 0.99 (0.067) |
| | (Q1,Q4) | 0.93 (0.068) |
| | (Q2,Q3) | 0.97 (0.063) |
| | (Q2,Q4) | 1.09 (0.074) |
| | (Q3,Q4) | 1.04 (0.074) |
| Diastolic blood pressure/Sex | (male,female) | 0.93 (0.031) |

Table S3: Sample sizes for siblings and unrelated sets. As described in **Fig. 3A**, for the comparison of prediction accuracies of polygenic scores based on standard and sib-GWAS, we first ascertain SNPs in a large sample of unrelated individuals (”Unrelated-discovery”) and then estimate the effect of these SNPs with a standard regression using unrelated individuals (”Unrelated-n*”) and, independently, using sib-regression (in the ”Siblings” set). Finally, we used the polygenic scores for prediction in a third sample of unrelated individuals (”Unrelated-prediction”). This table shows sample sizes used for each set across the traits analyzed.

| **Trait** | **Siblings (pairs)** | **Unrelateds-discovery** | **Unrelateds-n*** | **Unrelateds-prediction** |
|---|---|---|---|---|
| Age at first sex | 13677 | 244929 | 8843 | 27214 |
| Alcohol intake frequency | 17288 | 276872 | 10977 | 30763 |
| Basal metabolic rate | 16802 | 269728 | 13490 | 29969 |
| Birth weight | 6753 | 159237 | 5608 | 17693 |
| Body mass index | 17223 | 274887 | 12377 | 30543 |
| Diastolic blood pressure | 14795 | 253343 | 9424 | 28149 |
| Fluid intelligence | 3889 | 101070 | 2928 | 11229 |
| Forced vital capacity | 14611 | 252859 | 9731 | 28095 |
| Hair color | 16859 | 272151 | 11825 | 30238 |
| Hand grip strength | 17070 | 275067 | 10884 | 30563 |
| Height | 17248 | 270065 | 18085 | 30007 |
| Hip circumference | 17254 | 275957 | 11615 | 30661 |
| Household income | 13244 | 239274 | 8787 | 26585 |
| Neuroticism score | 11759 | 227111 | 6825 | 25234 |
| Overall health rating | 17195 | 276628 | 10365 | 30736 |
| Pack years of smoking | 2307 | 85626 | 1604 | 9513 |
| Pulse rate | 14791 | 253877 | 8790 | 28208 |
| Skin color | 16903 | 274150 | 10342 | 30461 |
| Waist circumference | 17257 | 275863 | 11757 | 30651 |
| Whole body fat mass | 16750 | 270569 | 12046 | 30063 |
| Whole body water mass | 16804 | 269694 | 13535 | 29965 |
| Years of schooling | 17041 | 273599 | 11873 | 30399 |
| Simulated trait 1 | 17305 | 276731 | 11362 | 30747 |
| Simulated trait 2 | 17305 | 276382 | 11749 | 30709 |

Table S4: Academic degree to years of schooling conversion table. The qualifications were converted to years of schooling following Okbay et al. [1]

| Qualifications (UKB data field 6138) | Years of schooling |
|---|---|
| College or University degree | 20 |
| NVQ or HND or HNC or equivalent | 19 |
| Other professional qualifications eg: nursing, teaching | 15 |
| A levels/AS levels or equivalent | 13 |
| O levels/GCSEs or equivalent | 10 |
| CSEs or equivalent | 10 |
| None of the above | 7 |

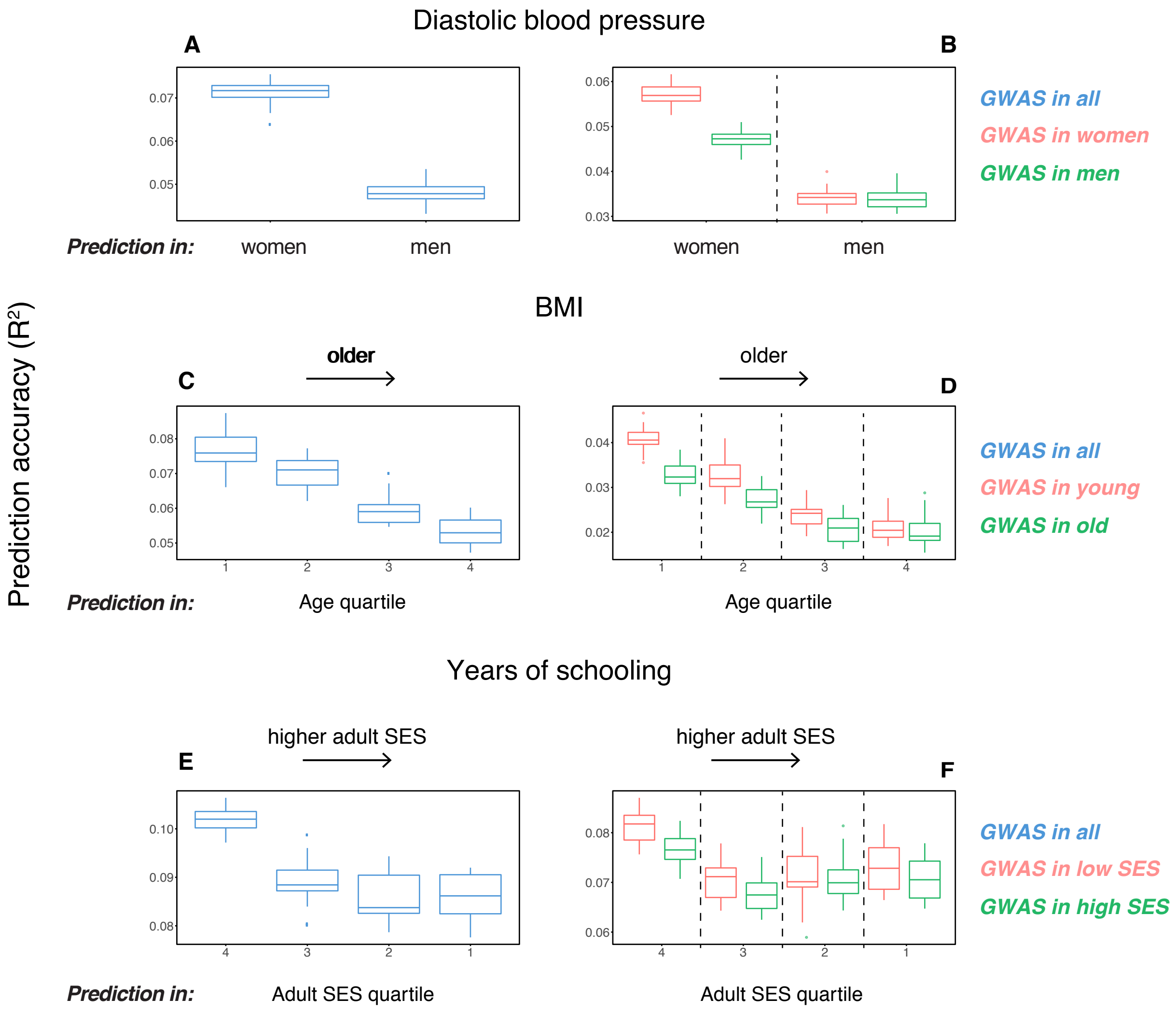

Figure S1: Variable prediction accuracy (measured as $R^2$) even within an ancestry group. This figure mirrors **Fig. 1** of the main text, except for the y-axis showing are $R^2$ values, rather than incremental $R^2$. Each box and whiskers plot was computed based on twenty choices of estimation and prediction sets. Thick horizontal lines denote the medians. **(A,C,E)** The polygenic scores were estimated in large samples of unrelated "White British" (WB) individuals. Phenotypes were then predicted in distinct sets of unrelated WB individuals, stratified by sex (A), age (C) or Townsend deprivation index, a measure of SES (E). **(B,D,F)** Same as in A,C,E, but here the polygenic scores are based on a GWAS in a sample limited to one sex or to one age or SES stratum. When the GWAS is performed in the sex or stratum that shows higher prediction accuracy in A,C,E (female, young, low SES), the qualitative trend is the same; but when the GWAS is performed in men, old or high SES groups, prediction accuracy is diminished and similar across prediction sets.

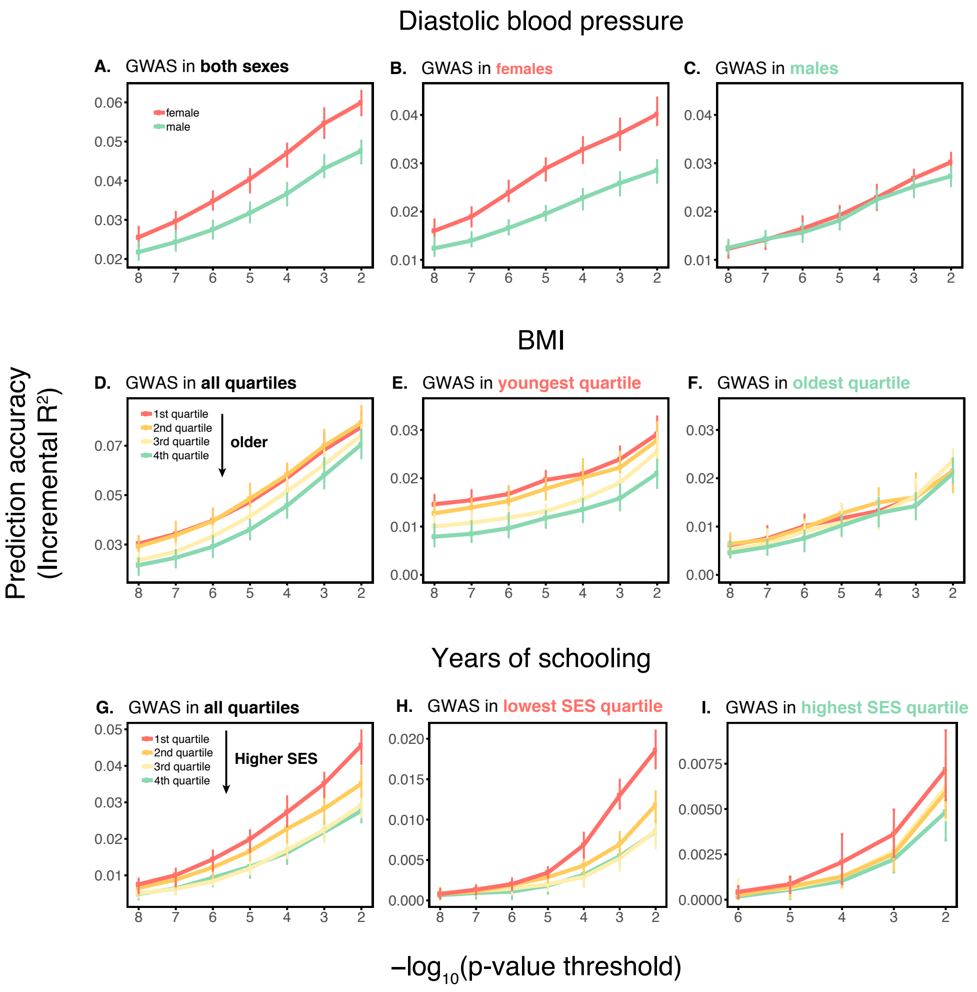

Figure S2: Dependence on the number of SNPs included in the polygenic score. This figure extends **Fig. 1** of the main text, showing the prediction accuracies as a function of the p-value threshold for inclusion of a SNP in the polygenic score. The higher the p-value threshold is, the more SNPs are included. Shown are incremental $R^2$ values in different prediction sets. Points and error bars are mean and central 80% range computed based on twenty choices of estimation and prediction sets. **(A,D,G)**. The polygenic scores were estimated in large samples of unrelated "White British" (WB) individuals. Phenotypes were then predicted in distinct samples of unrelated WB individuals, stratified by sex (A), age (D) or Townsend deprivation index, a measure of SES (G). **(B-I)** Same as in A,D,G, but here the polygenic scores are based on a GWAS in a sample limited to one sex, age or SES group. The trends shown in **Fig. 1** of the main text are for p-value threshold of $10^{-4}$, and are qualitatively similar to the trends at other p-value thresholds.

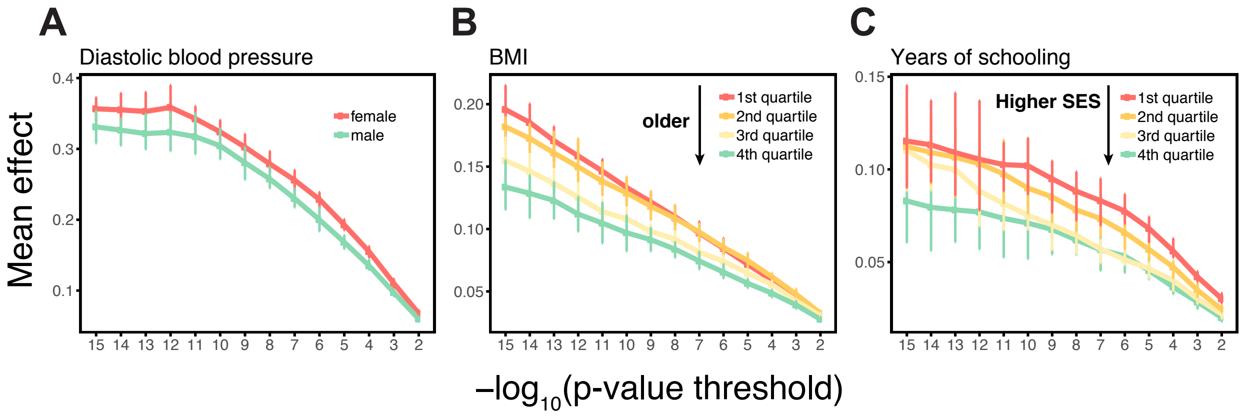

Figure S3: Estimating mean effect size across strata SNPs were ascertained in large samples of unrelated "White British" (WB) individuals. The effects of trait-increasing alleles were then re-estimated in an independent set of unrelated WB individuals (that were excluded from the original GWAS) stratified by sex for diastolic blood pressure **(A)**, by age for BMI **(B)** and by Townsend deprivation index, a measure of SES for years of schooling **(C)**. Points and error bars are mean and central 80% range computed based on twenty choices of ascertainment and estimation sets, plotted as a function of the p-value threshold (for p-values obtained in the original GWAS).

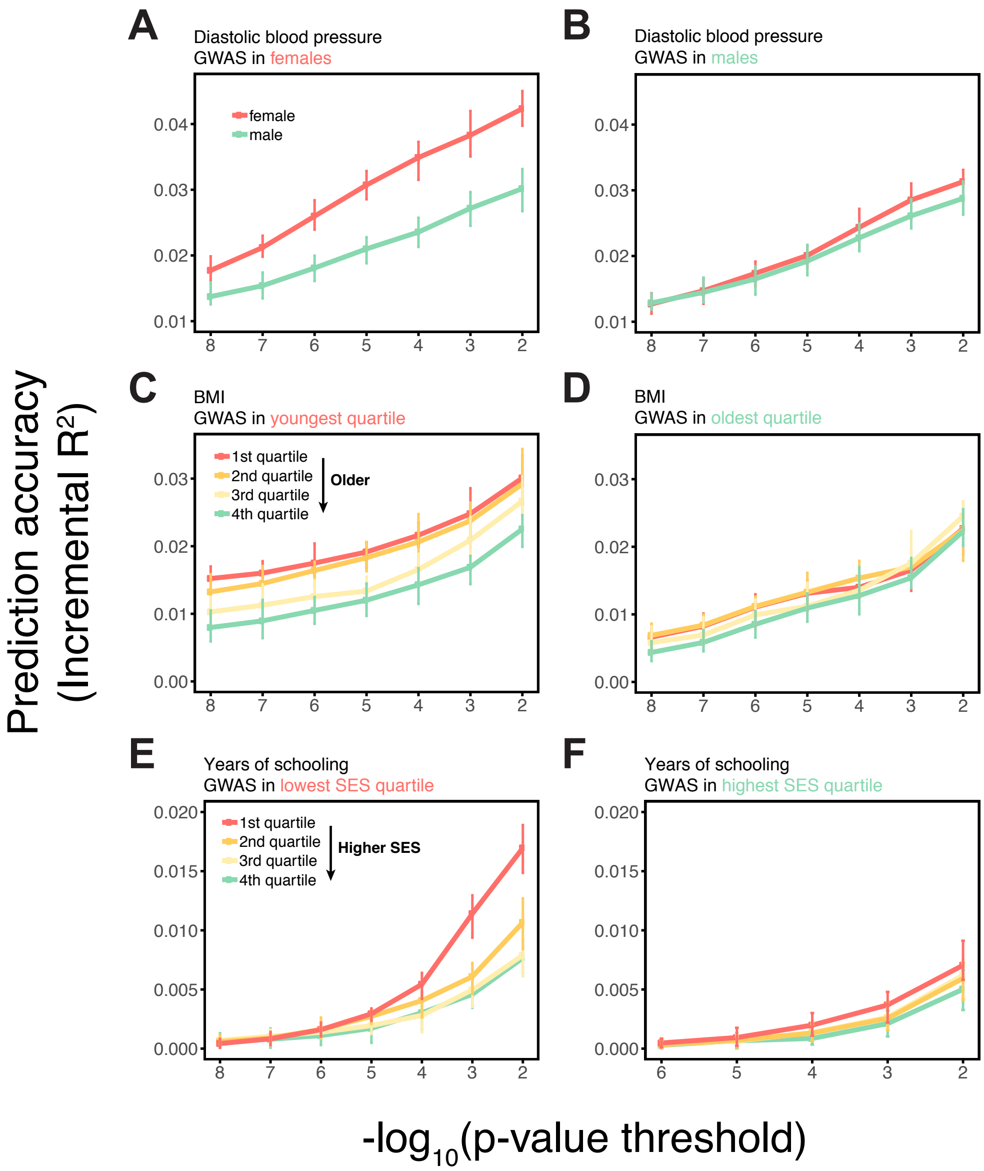

Figure S4: Variable prediction accuracy within an ancestry: sensitivity to controls for population structure. This figure mirrors the panels in **Fig. S3**, except that in these, the standard GWAS estimates were obtained from a linear mixed model (LMM). Shown are the prediction accuracies, measured as incremental $R^2$, as a function of the p-value threshold for inclusion of a SNP in the polygenic score. Points and error bars are mean and central 80% range computed based on twenty choices of estimation and prediction sets. The polygenic scores are based on a GWAS in a sample limited to one sex, age or SES group. Phenotypes are then predicted in distinct samples of unrelated individuals, stratified by sex **(A)**, age **(B)** or Townsend deprivation index, a measure of SES **(C)**. The qualitative trends are similar to those in **Fig. S3**, which uses a standard linear regression with PCs as a control for population structure when testing for an association between the phenotypes and genotypes. The similarity suggests that the observed differences in prediction accuracies across strata are not driven to a large degree by population structure confounding.

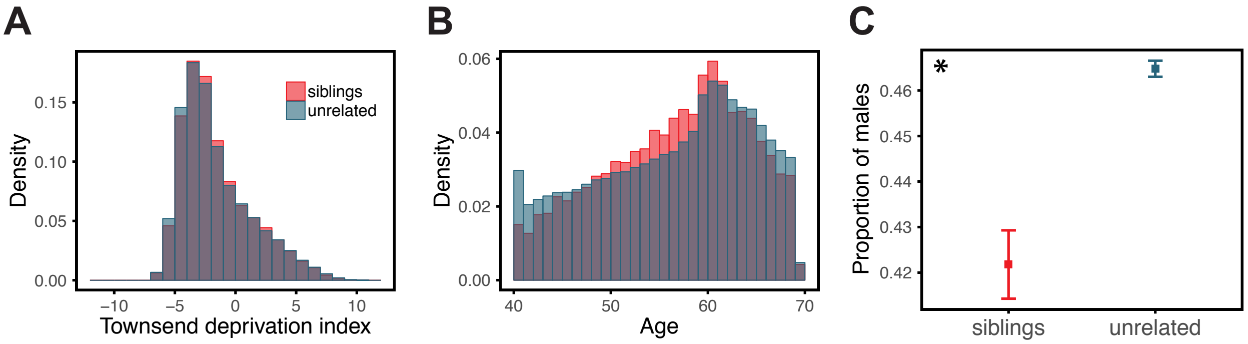

Figure S5: Comparison of siblings and unrelated individuals in the UK Biobank with respect to age, SES, and sex ratio. Panels show the distribution of Townsend deprivation index, a measure of SES **(A)**, the age distribution **(B)**, and the proportion of males **(C)** for the siblings and unrelated sets used in the analysis described for **Fig. 3** of the main text. For each sibling pair, one sibling was randomly selected for these comparisons. The asterisk symbol marks a significant difference between siblings and unrelated individuals, Mann-Whitney $p < 0.01$. SES and age distributions are quite similar in siblings and unrelated sets, whereas the proportion of males is slightly but significantly smaller in the siblings.

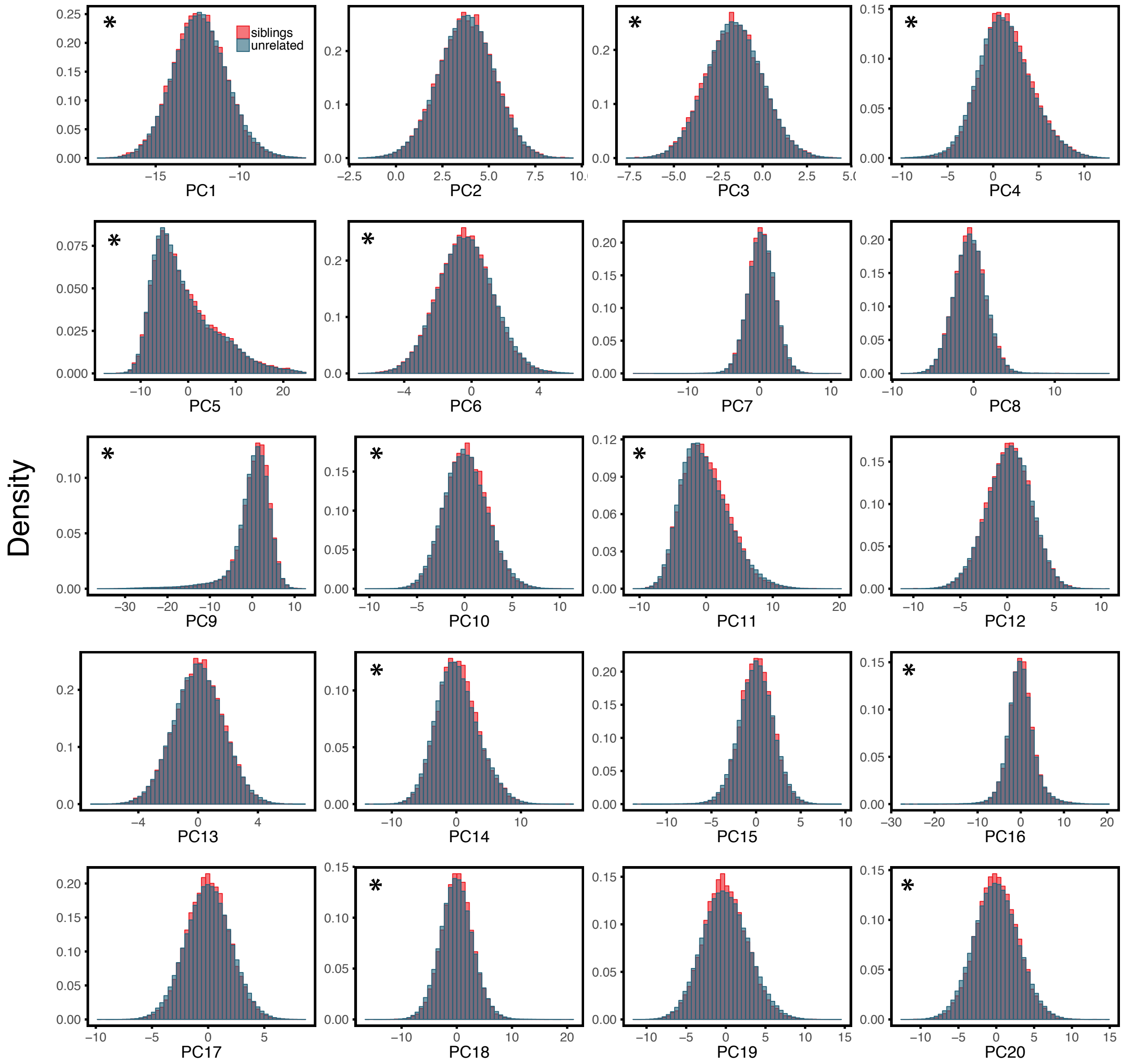

Figure S6: Comparison of siblings and unrelated individuals in the UK Biobank with respect to population structure. Panels show the distribution of PCs for the siblings and unrelated sets used in the analysis described for **Fig. 3** of the main text. For each sibling pair, one sibling was randomly selected for these comparisons. The asterisk symbol marks a significant difference between siblings and unrelated individuals, Mann-Whitney $p < 0.01$. Siblings and unrelated sets are broadly similar with respect to their genetic ancestries.

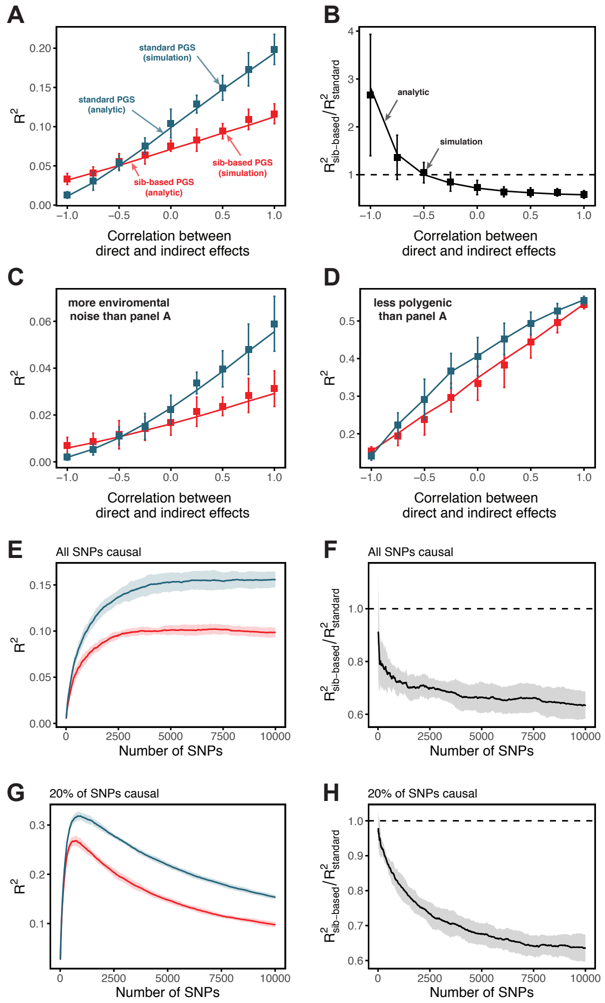

Figure S7: Simulation results for polygenic scores based on standard GWAS and sib-GWAS in the presence of indirect effects. **(A,B)** Simulation results as a function of the correlation between direct and indirect effects, $\rho$. Simulations were performed with $h^2_\beta = 0.5$, $\tau^2_\eta = 0.1$, and $\sigma^2_\beta/\sigma^2_\eta = 5$. The size of the estimation set in the sib-GWAS is 10,000, and the size of the estimation set in the standard GWAS is chosen to match sampling variances between the two study designs. The polygenic scores is based on 10,000 causal loci; its performance was evaluated in an independent set of 10,000 unrelated individuals. As long as direct and indirect effects are not strongly negatively correlated, out-of-sample prediction accuracy is higher for the polygenic scores based on standard GWAS. **(C)** Same as **(A)** but with three-fold greater environmental noise. **(D)** Same as (A) but with 100 causal loci. In (A-D) points are mean $\pm$ 2 SD in 10 simulation iterations. Solid lines are values based on analytic expressions derived in Section 1.3.2. **(E-H)** Simulations results, with the same parameters as in (A) but $\rho = 0.5$, as a function of the number of SNPs included in the polygenic scores, with all loci being causal (E,F), or with 20% of loci being causal (G,H). SNPs are added in an increasing order of their association p-value in an independent set of 20,000 unrelated individuals. In both cases, the ratio of prediction accuracies of polygenic scores based on sib- versus standard GWAS becomes smaller with the inclusion of more weakly associated SNPs, a behavior qualitatively similar to observations in Fig. 3 in the main text. Points are mean $\pm$ 2 SD in 10 simulations.

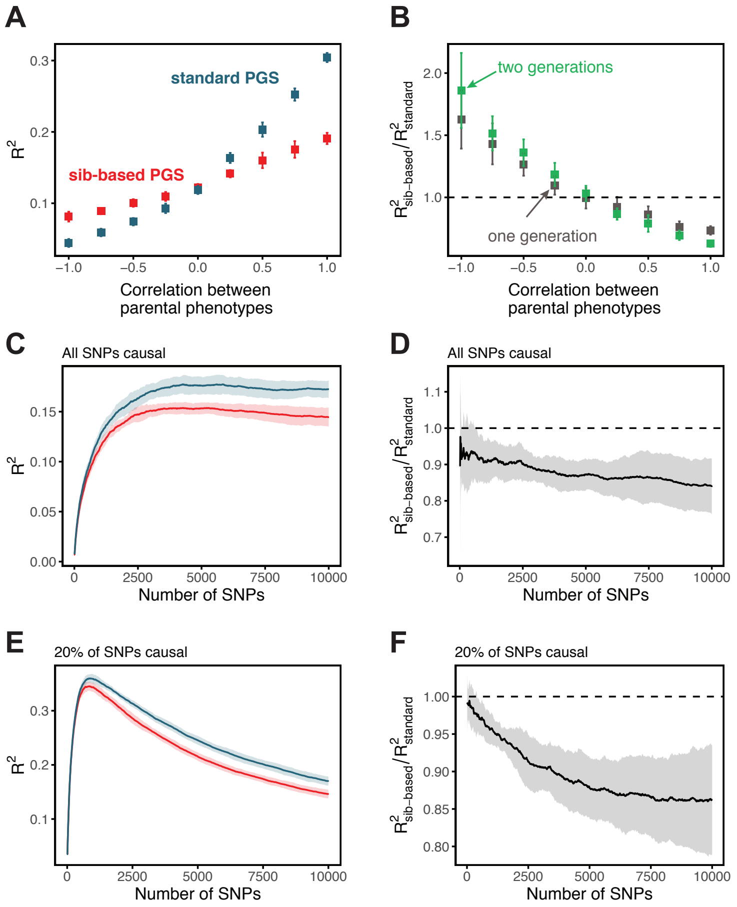

Figure S8: Simulation results for polygenic scores based on standard GWAS and sib-GWAS in the presence of assortative mating. **(A)** Simulation results as a function of the approximate correlation between parental phenotypes, $\rho_a$. Simulations were performed with $\tau^2 = 0.5$ under random mating. The size of the estimation set in the sib-GWAS is 10,000, and the size of the estimation set in the standard GWAS is chosen to match sampling variances between the two study designs. The polygenic scores is based on 10,000 causal loci; its performance was evaluated in an independent set of 10,000 unrelated individuals. Standard-GWAS based polygenic scores outperforms (underperforms) sib-GWAS based polygenic scores under positive (negative) assortative mating. **(B)** Ratio of prediction accuracies of the polygenic scores based on sib- versus standard GWAS, as a function of $\rho_a$, for two sets of simulations with one or two generations of assortative mating, with same parameters as in (A). **(C-F)** Simulations results, with the same parameters as in (A) but $\rho_a = 0.5$, as a function of the number of SNPs included in the polygenic score, with all loci being causal (C,D), or with 20% of loci being causal (E,F). SNPs are added in the order of their association p-value in an independent set of 20,000 unrelated individuals. In both cases, the ratio of prediction accuracies for scores based on sib-GWAS versus standard GWAS becomes smaller with the inclusion of more weakly associated SNPs, a behavior that is qualitatively similar to observations in **Fig. 3** in the main text. Points are mean ± 2 SD in 10 simulation iterations.

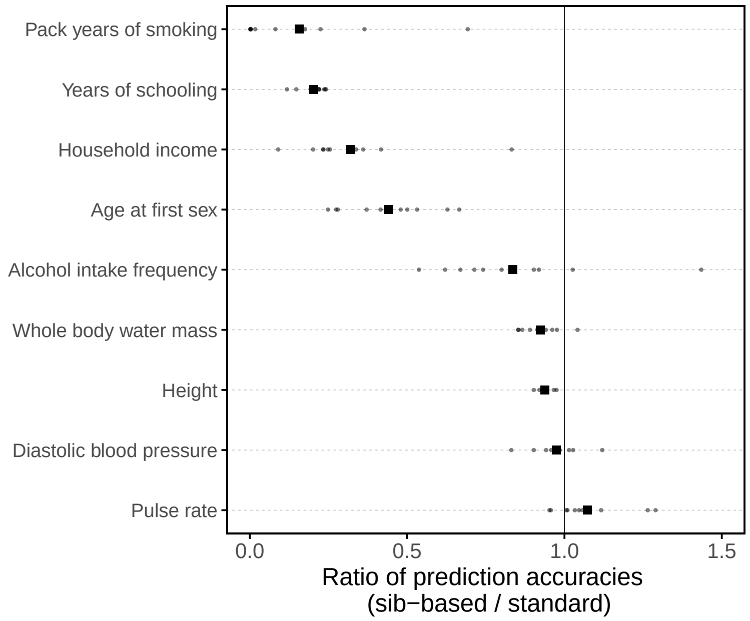

Figure S9: Comparison of prediction accuracies of polygenic scores based on standard and sib-GWAS matched for sex ratio. This figure mirrors **Fig. 3B** of the main text, but here the samples of siblings and unrelated individuals used in the analysis are matched for sex ratio. Results are shown for diastolic blood pressure, as the prediction accuracy differed between sexes (**Fig. 1**), the related phenotype of pulse rate, and a subset of the traits for which the prediction accuracy varied by GWAS design (**Fig. 3B**). Small points show the ratio of the prediction accuracies in the two designs across 10 iterations; in each iteration, we resample sets of unrelated individuals to constitute three sets: for discovery, estimation and prediction. Large points show mean values.

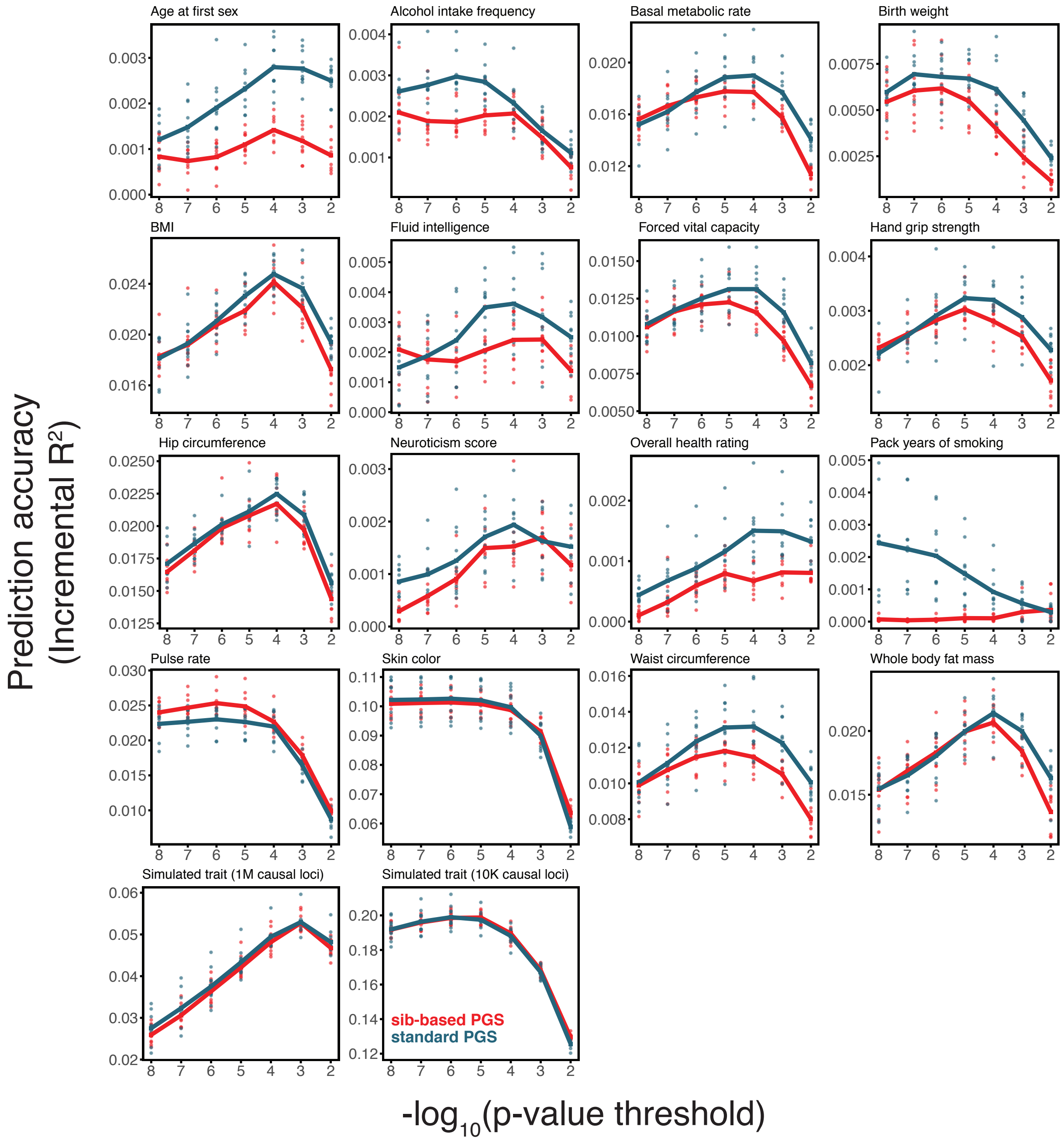

Figure S10: Prediction accuracy of polygenic scores based on sib-and standard GWAS, for a range of traits. This figure complements **Figs. 3C-H** of the main text, showing the results of the study design depicted in **Fig. 3A** for all traits presented in **Fig. 3**. As described for **Fig. 3**, we randomly divided unrelated individuals to constitute three non-overlapping sets for discovery, estimation and prediction. Small points correspond to 10 iterations of resmapling these three sets. The prediction accuracy is plotted as a function of the p-value threshold, where p-values come from the discovery GWAS. Lines show mean values.

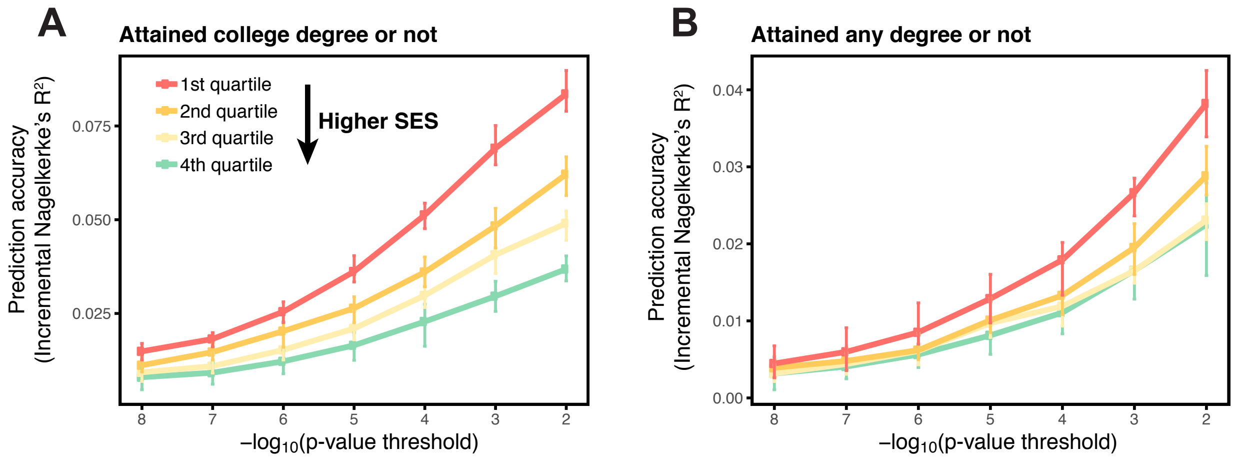

Figure S11: Variable prediction accuracy within an ancestry group for binary education phenotypes. Shown are incremental Nagelkerke's $R^2$ for two binary education phenotypes, attained a college degree or not **(A)**, and attained any degree or not **(B)**. The polygenic scores are estimated in large samples of unrelated individuals. Phenotypes are then predicted in distinct samples of unrelated individuals, stratified by Townsend deprivation index, a measure of SES. Points and error bars are mean and central 80% range computed based on twenty choices of estimation and prediction sets, plotted as a function of the p-value threshold (with p-values obtained in the original GWAS).

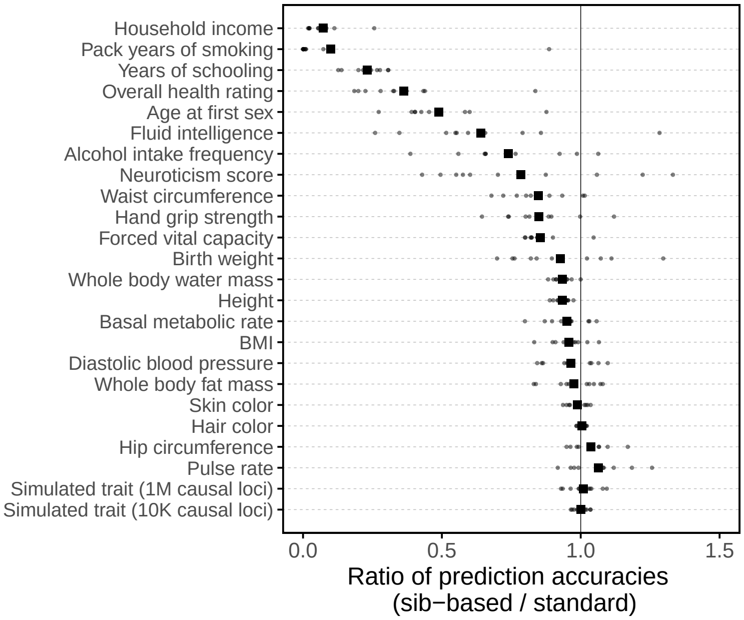

Figure S12: Comparison of prediction accuracies of polygenic scores (measured as $R^2$) based on standard and sib-GWAS. This figure mirrors **Fig. 3B** of the main text, but here we first residualized the phenotypes on covariates, and then ran the same pipeline described as that used to generate **Fig. 3B** on the residuals without further adjustment for covariates in the GWAS or prediction evaluation. Thus, this figure relates more directly to the analytical derivation in **Section 1.2**. However, the results in **Fig. 3B** and here are qualitatively similar. Small points show the ratio of the prediction accuracies in the two designs across 10 iterations; in each iteration, we resample sets of unrelated individuals to constitute three sets: for discovery, estimation and prediction. Large points show mean values.
